# Plot diversity differentially affects the chemical composition of leaves, roots and root exudates in four subtropical tree species

**DOI:** 10.1101/2020.11.26.400424

**Authors:** Alexander Weinhold, Stefanie Döll, Min Liu, Andreas Schedl, Xingliang Xu, Steffen Neumann, Nicole M. van Dam

**Affiliations:** German Centre for Integrative Biodiversity Research (iDiv) Halle-Jena-Leipzig, Puschstraße 4, 04103 Leipzig, Germany; Institute of Biodiversity, Friedrich Schiller University Jena, Dornburger-Str. 159, 07743 Jena, Germany; Key Laboratory of Ecosystem Network Observation and Modeling, Institute of Geographic Sciences and Natural Resources Research, Chinese Academy of Sciences, 11A, Datun Road, Chaoyang District, 100101 Beijing, China; Synergy Research Group Bioinformatics and Scientific Data (BASDA), Leibniz Institute of Plant Biochemistry, Weinberg 3, 06120 Halle, Germany; College of Resources and Environment, University of Chinese Academy of Sciences, Yanqi Lake, Huairou District, 101408 Beijing, China

**Keywords:** Chemical diversity, ecometabolomics, LC-qToF-MS, plant-plant interactions, root exudate, secondary metabolites, tree interactions, tropical forest ecology

## Abstract

1. Plants produce thousands of compounds, collectively called the metabolome, which mediate interactions with other organisms. The metabolome of an individual plant may change according to the number and nature of these interactions. We tested the hypothesis that tree diversity level affects the metabolome of four subtropical tree species in a biodiversity ecosystem-functioning experiment, BEF-China. We postulated that the chemical diversity of leaves, roots and root exudates increases with tree diversity. We expected the strength of this diversity effect to differ among leaf, root and root exudates samples. Considering their role in plant competition, we expected to find the strongest effects in root exudates.
2. In an ecometabolomics approach, roots, root exudates and leaves of four tree species *(Cinnamomum camphora, Cyclobalanopsis glauca, Daphniphyllum oldhamii, Schima superba)* were sampled from selected plots in BEF-China. Samples were extracted and analysed using Liquid Chromatography-Time of Flight-Mass Spectrometry. The exudate metabolomes were normalized over their non-purgeable organic carbon level. Multivariate analyses were applied to identify the effect of both neighbouring (local) trees and plot diversity on tree metabolomes. The species and sample specific metabolites were assigned to major compound classes using the ClassyFire tool, whereas m/z features related to diversity effects were annotated manually.
3. Individual tree species showed distinct leaf, root and root exudate metabolomes. The main compound class in leaves were the flavonoids, whereas carboxylic acids, prenol lipids and specific alkaloids were most prominent in root exudates and roots. Overall plot diversity had a stronger effect on metabolome profiles than the diversity of local, directly neighbouring trees. Leaf metabolomes responded more often to tree diversity level than exudates, whereas root metabolomes varied the least. We found not overall correlation between metabolite richness or diversity and tree diversity.
4. Synthesis: Classification of metabolites supported initial ecological interpretation of differences among species and organs. Particularly the metabolomes of leaves and root exudates respond to differences in tree diversity. These responses were neither linear nor uniform and individual metabolites showed different dynamics. More controlled interaction experiments are needed to dissect the causes and consequences of the observed shifts in plant metabolomes.

## INTRODUCTION

Forests are important ecosystems that provide many ecosystem services. To humans, they are an important resource for food, fire wood and timber (Jang et al., 2019). In addition, forests harbour an enormous number of different organisms, from microbes to large mammals (Barlow et al., 2018). This is particularly true for (sub) tropical forests, which contain 96% all tree species on earth (Poorter et al., 2015). Consequently, tropical forests sustain high levels of biodiversity, in particular insects and other arthropods. A study in Panama estimated that one hectare of subtropical forest contains 60% of insect diversity on the larger landscape scale (Basset et al., 2012). Last, but not least, forests may mitigate the effects of climate change (Trumbore et al., 2015). In particular, tropical forests can reduce warming through evaporative cooling (Bonan, 2008). Forests also sequester large amounts of carbon, thereby storing about 45% of total terrestrial C (Bonan, 2008). Tree species diversity commonly has a positive effect on these ecosystem services. Several studies show positive relationships between tree species-richness and insect diversity (Schuldt et al., 2019), biomass production (Guillemot et al., 2020), and carbon storage (Poorter et al., 2015, Steur et al., 2020). Finally, trees in species-diverse forests may reduce drought and heat stress by reducing forest floor temperature. The structural diversity of trees in more diverse forests enhances canopy closure, thereby mitigating temperature effects on organisms on the forest floor (Guillemot et al., 2020). Considering the current biodiversity crisis fuelled by anthropogenic changes in land use and CO_2_ levels (Trumbore et al., 2015), it is important to understand the underlying mechanisms driving diversity in forests.

The positive relationships between tree diversity and ecosystem services are for a part ascribed to complementarity effects. Complementarity can emerge on three different levels: resource partitioning, abiotic facilitation, and biotic feedbacks (Barry et al., 2019). Resource partitioning between plant species over chemical, temporal, or spatial scales promotes the coexistence of species that compete for limited resources (Tilman, 1982, McKane et al., 2002). For example, tropical tree species can enhance light capture via crown plasticity and spatial and temporal niche differentiation (Sapijanskas et al., 2014). At the same time, they can optimize the use of soil resources through root plasticity (Sun et al., 2017) and mycorrhizal networks to reduce competition (Schenk, 2006, Simard and Durall, 2004). Abiotic facilitation may occur when a species benefits from being in a more diverse community, e.g. when shallow-rooting tree species suffer less from drought stress (Fichtner et al., 2020, Vitali et al., 2018). Biotic feedbacks may contribute to positive density dependent effects, for example, when trees share mycorrhiza and generalist pollinators. Interactions with specialist pathogens and herbivores may lead to negative density dependent effects. Tree performance in monocultures would be reduced compared to the same species in mixed communities, due to the dilution of specific enemies in the latter (Barry et al., 2019). Plant compounds play an important role in interactions between plants and their biotic environment. Plant secondary metabolites, such as alkaloids, phenolic compounds or volatile organic compounds, mediate resistance to herbivores and pathogens, attract pollinators and beneficial root microbes, and serve as allopathic compounds to suppress competing plant species (Raguso et al., 2015). In contrast to primary compounds, such as amino acids and sugars, the profiles of secondary metabolites are species specific (Sedio et al., 2017, Endara et al., 2015, Sardans et al., 2020). Metabolomics is principally an untargeted technology and can be applied to any species without prior knowledge of its chemical composition (Macel and van Dam, 2018). This independence also allows the application of metabolomics in ecology, which gave rise to the field of ecometabolomics (Peters et al., 2018, Sardans et al., 2020). Ecometabolomic approaches enhance our understanding of the role of plant metabolites in individual plant interactions as well as in ecosystem functioning, for example as a functional trait in plant diversity (*Walker et al., editorial of this issue*).

Metabolomic analyses of tropical tree leaf metabolomes showed that there is substantial variation in secondary metabolite profiles among (congeneric) species (Endara et al., 2015, Sedio et al., 2017). The authors hypothesized that this chemical diversity has arisen from selection pressures exerted by enemies shared among sympatric congeners, which has led to chemical diversification within several speciose tropical genera (Endara et al., 2015, Richards et al., 2015, Sedio et al., 2017). However, plant metabolomes are not static, and change in response to biotic and abiotic environmental variation. For example, attacks by aboveground or belowground herbivores and pathogens may shift the metabolome of plants towards producing more (defensive) metabolites (Macel et al., 2014, van Dam and Heil, 2011). Variation in abiotic environmental factors, such as temperature or water availability, strongly affects plant physiological processes and thereby plant metabolomes (Sardans et al., 2020, Walker et al., 2019). In other terms, plant metabolomes are plastic and change depending on the type and number of interactions in their environment.

Ecometabolomic analyses in biodiversity and ecosystem functioning (BEF) studies manipulating diversity of grassland species showed that primary and secondary metabolomes of individual plant species change with plant diversity level (Huberty et al., 2020, Ristok et al., 2019, Scherling et al., 2010). These diversity-driven effects on plant metabolomes could be caused by plant-plant interactions or soil feedback. Plant-plant interactions may cause facilitation as visible in increased amino acid levels (Scherling et al., 2010) or competition, leading to decreased levels of amino acid levels (Scherling et al., 2010) or the production of allelopathic metabolites (Fernandez et al., 2016). Soil feedbacks occur because plant species and plant communities facilitate specific soil microbial communities (Bardgett and van der Putten, 2014). Individual plants exposed to these different soil communities can respond by changing their metabolomic profiles (Huberty et al., 2020, Ristok et al., 2019). Similar effects were observed on trees using targeted metabolomic approaches. For example, total phenolic concentrations in birch (*Betula pendula*) leaves increased with plot diversity level (Poeydebat et al., 2020). So far, untargeted metabolomic approaches have been mostly used to quantify chemical diversity related to phylogenetic distances in large tropical tree genera (Endara et al., 2015, Salazar et al., 2018, Sedio, 2017), chemical diversity within canopies (Sedio et al., 2019, Sedio et al., 2017, Wiggins et al., 2016), induced responses, or variation in leaf metabolomes among seasons (Gargallo-Garriga et al., 2020). To our knowledge, none of these studies considered the effect of tree diversity levels as a driver of variation in tree metabolomes.

To test whether tree metabolomes are affected by diversity levels in their surroundings, we analysed root exudate, root and leaf metabolomes of four subtropical plant species in a biodiversity and ecosystem functioning (BEF) experimental field site in China (Trogisch et al. unpublished data) (Bruelheide et al., 2014). Whereas other studies on tree metabolomes focus on leaf metabolomes, we explicitly included analyses of roots and their exudates. This is important, because roots and shoots have different functions and function in a different biotic and abiotic environments (van Dam, 2009). Roots fulfil important physiological functions, such as water uptake, macro- and micro-nutrient foraging and storage. Niche differentiation, in the form of morphological root responses (Sun et al., 2017) and the uptake of different forms of N may alleviate nutrient imitation and maintain tree diversity (Lang et al., 2014). Moreover, roots produce root exudates which are secreted into the rhizosphere and contain a large variety of metabolites (van Dam and Bouwmeester, 2016). In nutrient-poor soils, the chemicals in root exudates increase mineral nutrient bioavailability through rhizosphere priming effects (Dijkstra et al., 2013) and mediate mutualistic interactions with beneficial microorganisms, such as mycorrhizal fungi (Ferlian et al., 2018). In addition, they deter or resist pathogenic microbes, invertebrate herbivores and parasitic plants, as well as suppress competing plant species (Baetz and Martinoia, 2014, Zeng, 2014). In consequence, root exudates might play a key role in direct and indirect tree-tree interactions. We therefore expect that both root, root exudate and leaf metabolomes respond to tree diversity levels in their environment.

Root exudates, root and leaves of the four selected tree species were sampled in plots with different levels of diversity, ranging from monocultures to 16 and 24 species plots. We also recorded the local species diversity among the trees directly surrounding the target trees. This allowed us to test the effect of surrounding tree diversity levels (local diversity) and total plot diversity on the metabolomes of leaves, roots and root exudates. We hypothesized that roots, root exudates and leaves showed species-specific metabolomic profiles, and that within species, root and root metabolomes would be most similar. We also expected that local neighbour diversity would have a larger effect on plant metabolomes than plot level diversity, and that metabolite diversity would increase with tree diversity level. Finally, we postulated that the metabolomes of root exudates respond stronger to plot diversity than roots or leaves, because of the positive effects of diversity on the production of fine roots (Sun et al., 2017), which might also enhance root exudation.

## MATERIALS AND METHODS

### Field location

This study was carried out in the Biodiversity–Ecosystem Functioning Experiment China (BEF-China), which has been set-up and managed since 2008 (Bruelheide et al., 2014). It is located in Jiangxi Province, China (29°08′–29°11′N, 117°90′–117°93′E) on two sites, A and B, which were planted in 2009 and 2010, respectively. A former timber forest was replanted with local tree species in monoculture and mixed stands using a “broken stick” design. Using a pool of 40 tree species, extinction scenarios were simulated with tree richness levels of 1, 2, 4, 8, 16 and 24 species on a total of 566 plots of 25.8 m × 25.8 m and 400 trees each. The trees are planted on a rectangular grid with 1.29 m distance. Thus, every tree has eight potential neighbours at a distance of 1.29 m or 1.82 m (diagonal).

This study was carried out on site B, which ranges in altitude from 113 to 182 m and with slopes from 15–43 degrees. The site has a subtropical climate. From 1971–2000 the mean annual temperature was 16.7 °C and the mean annual precipitation 1800 mm. This has increased to 17.9 °C and 2076 mm, respectively, during 2013–2017. January is the coldest month with a mean temperature of 0.4 °C and July is the hottest with a mean temperature of 34.2 °C. The (natural) vegetation is characterized by subtropical forest with a mixture of evergreen and deciduous species.

### Target tree species

For our analyses, we targeted four different tree species planted in BEF China Site B (Fig. S1). In 2018, 8 tree species pairs (TSP) consisting of 16 tree species were labelled for various diversity (1, 2, 4, 8, 16/24) analyses in the International training Network TreeDì (Zhang et al., 2020). In this study, we selected four of these species that had sufficient replicates (3 or more) in plots of different diversity levels. Moreover, we selected trees from different families to avoid phylogenetic bias.

*Cyclobalanopsis glauca* (Thunb.) Oerst., (homotypic syn. of *Quercus glauca* Thunb.), Fagaceae, or the “ring-cup oak”, is native to subtropical and warmer temperate zones of Asia. It contains a variety of tannins (catechins, procyanidins, gallic acids), triterpenoids, flavonoid glycosides and steroids (cycloartanols, stigmastanes) (Kamano et al., 1976, Shen et al., 2012, Sheu et al., 1992, Suga and Kondo, 1974, Wakamatsu et al., 2020)). *Schima superba* Gardner & Champ., Theaceae, is a dominant broad-leaved evergreen tree distributed over temperate Asia. Chemical analyses showed that *Schima* spp. contain triterpenoids and saponins as well as flavonoids and anthocyanins (Deng et al., 2010, Kitagawa et al., 1975, Liang et al., 2019, Liu et al., 2019, Wu et al., 2019, Wu et al., 2015, Yang et al., 2018, Yu et al., 2019). *Daphniphyllum oldhamii* (Hemsl.) K. Rosenthal, heterotypic synonym of *Daphniphyllum pentandrum* Hayata, Daphniphyllaceae, is native to temperate Eastern Asia. Species in the genus *Daphniphyllum* contain Daphniphyllum alkaloids, a diverse group of specific alkaloids with unusual ring structures (azaspirodecane derivatives) with different backbones, probably derived from squalene and mevalonate (Kobayashi and Kubota, 2009). Furthermore, lignans and flavonoids have been described (Chao et al., 2018, Gan et al., 2007, Kobayashi et al., 2003, Mu et al., 2006, Mu et al., 2007, Shao et al., 2004, Takatsu et al., 2004). *Cinnamomum camphora* (L.) J. Presl, camphor, Lauraceae, is native to Japan and Taiwan and cultivated or naturalized in temperate and tropical regions worldwide. This plant is well known for its monoterpenoids (bicyclic, acyclic, menthane monoterpenoids, e.g. camphor, camphene, terpineol, pinene, and borneol). *C. amphora also* contains tri- and sesquiterpenoids, neolignans, steroids, benzenoids, flavonoids and oxolanes (KNApSAcK database (Afendi et al., 2012)).

### Sampling design

The sampling took place from 19^th^ of October until 28^th^ of October 2019. We took our samples in the 13 plots in the BEF China experiment that contained our target species. We sampled root exudates, roots and leaves of the above-mentioned four tree species in plots with species richness levels 1, 2, 4, 8, or 16(24) species. In monoculture plots (1 per species), we sampled three randomly selected trees. In 2-species plots, 4-8 replicates were sampled for each species. In plots with 4, 8, 16/24 species, we sampled 2-7 trees per species. The detailed sampled species combinations and sampling replicate numbers were shown in Fig S1. In total, we sampled 84 trees, yielding 84 samples for root exudates, roots and leaves each.

After scrutiny of the metabolomes, it became apparent that 9 of the 84 root (exudate) samples were not from the target tree. Pre-experiments showed that under field conditions, the root excavated for exudate collection is not always originating from the target tree, even when taking all possible care. Eight of the root and root exudate samples could be reassigned to one of the other three species based on their metabolome. One sample could not be reassigned, thus, for the roots, 83 samples remained for statistical analyses, for the exudates 81 (2 were lost during sample preparation) and for the leaves the complete 84.

### Leaves, roots, and root exudate collection

For the collection of root exudates, we followed the protocol of (Phillips et al., 2008), with slight modifications. The root exudates were collected from the middle of two trees under each combination (Fig. S1). We carefully excavated the roots from the upper 10 cm of mineral soil, starting from the base of the tree, to identify terminal fine root strands (< 2 mm). Adhering soil particles were carefully removed with demineralized water. Roots were gently dried using paper tissue and placed into a 30 ml glass syringe, which was then filled with new, unused glass beads (500-750 μm, acid washed, ACROS Organics, ThermoFisher, New Jersey, USA/Geel, Belgium) The outlet of the glass syringe was attached to a plastic tube with a plastic 50 ML syringe attached to it. The glass syringe with the roots was filled with nutrient solution (0.50 mM NH_4_NO_3_, 0.10 mM KH_2_PO_4_, 0.20 mM K_2_SO_4_, 0.15 mM MgSO_4_, 0.30 mM CaCl_2_) from the top. The solution was sucked through the plastic syringe and discarded. The root was left to equilibrate for 20 min, after which fresh nutrient solution was added and the procedure was repeated. After this second washing step, the glass syringe was closed with cotton wool and sealed with parafilm. Everything was wrapped in aluminium foil and left for 48 h. After 48 h, the roots were cut off and the syringes, which contained the root and glass beads, were placed in plastic bags and brought into the lab. In addition, randomly chosen leaves were sampled for each tree from the shaded part of the crowns. Each target tree’s position and neighbourhood species were recorded at the time of sampling. These data were used to calculate neighbourhood diversity i.e. the number of species that were directly neighbouring the target tree.

To recover the root exudates, a 0.22 μm sterile filter was placed on a vacuum pump manifold over a 60 mL glass/plastic vial. The glass syringe with the roots and the glass beads was mounted on top of the manifold and filled with 60 ML nutrient solution. Then, the vacuum pump was started and the exudate solution was collected into the vial. Five ml of the exudate solution of each sample was aliquoted for Non-Purgeable Organic Carbon (NPOC) analysis (see below). The remaining root exudate solution was frozen at −20 °C in 60 ml HDPE cryo-vials until freeze-drying but thawed for shipment. After the collection of exudate solution, the roots were cleaned with water to remove residual glass beads. Root and leaf samples were put in envelopes, oven dried for 30 min under 105 °C, and then oven dried for 24 h under 60 °C.

### Sample preparation for LC-MS

Dried roots and leaves were extracted for LC-MS according to a standard protocol: Per 20 mg powdered material, 1 ml of extraction buffer (75% v/v methanol, HPLC grade, 25% v/v acetate buffer (2.3 mL acetic acid and 3.41g ammonium acetate in 1L 18MΩ water, pH set to 4.8) plus 50µl 100mM IAA-Valin as internal standard) was added and shaken with ceramic beads in a tissue homogenizer (Retch MM400, Retch GmbH, Haan, Germany) for 5 min at 30Hz. Samples were centrifuged for 15 min at RT at 15.000g. The pellet was reextracted with another 1 ml of extraction buffer per 20 mg starting material, centrifuged again and both supernatants were unified. Samples were diluted 1:5 with the extraction buffer, kept at −20°C overnight, centrifuged at 15.000 g for 10 min and transferred to HPLC vials. Exudates were freeze-dried after shipment, redissolved in 1 ml water and the bottles washed with 1 ml methanol. The samples were transferred to a new 2 ml reaction tube, completely dried (vacuum centrifuge 27°C) and we tried to redissolve them in a smaller volume of the extraction buffer. However, this was not possible, so 1 ml of water was added, dissolution was aided by vortexing, ultrasonic bath and heating to 40°C, the samples centrifuged, the remaining pellet again dissolved in another 1 ml of water, centrifuged again and both supernatants were unified. After that, the samples were concentrated, but not to complete dryness, in a vacuum centrifuge.

### NPOC measurements

Non-purgeable organic carbon (NPOC) was determined after removal of inorganic C (acidification and sparging) by catalytic thermal oxidation (at 680°C) and subsequent detection of CO_2_ by an infrared gas analyser (TOC VCPN, Shimadzu, Germany) on three analytical replicates.

### Normalization of exudate samples by NPOC before LC-MS analysis

Because exudate NPOC values depended heavily on the root morphology and differed significantly among species (Fig. S2) we used NPOC values to dilute samples before analysis. This procedure normalizes for exudation rates and thus allows a better comparison of root exudation profiles over species and treatments. To correct for the NPOC, the volume of the concentrated samples was measured by taking it up with a pipette. The median NPOC of all samples was calculated (5.023 mg*L^−1^). The minimum was 0.84 mg*L^−1,^ the maximum was 33.66 mg*L^−1^. It was decided that the median concentration of carbon should correspond to 300μl, and the required volume for each sample was calculated as follows: volume _sample_ [μl] = NPOC _sample_ [mg.L^−1^]*300μl/ median NPOC [mg.L^−1^]. The concentrated samples (remaining volume 50-200 μl) were filled up with 70% methanol to attain the calculated volume (range 50 - 2011μl).

### LC-MS measurements

Chromatographic separations were performed at 40°C on an UltiMate™ 3000 Standard Ultra-High-Performance Liquid Chromatography system (UHPLC, Thermo Scientific) equipped with an Acclaim^®^ Rapid Separation Liquid Chromatography (RSLC) 120 column (150 × 2.1 mm, particle size 2.2 μm, ThermoFischer Scientific) using the following gradient at a flow rate of 0.4 ml min−1: 0–1 min, isocratic 95% A [water/formic acid 99.9/0.1 (v/v %)], 5% B [acetonitrile/formic acid 99.9/0.1 (v/v %)]; 1-2 min, linear from 5 to 20% B; 3–8 min, linear from 20 to 25% B; 8–16 min, linear from 25 to 95% B; 16-18 min, isocratic 95% B; 18 −18.01 min, linear from 95 to 5% B; 18.01-20 min, isocratic 5% B. Data was recorded from 0 min to 18 min. The injection volume was 10 μl for exudates and 5 μl for roots and leaves.

Eluted compounds were detected from m/z 90 to 1600 at a spectra rate of 5 Hz, (line spectra only) using an ESI-UHR-Q-ToF-MS (maXis impact, Bruker Daltonics) in positive ion mode with data dependent collision induced dissociation (Auto-MSMS mode). The following instrument settings were applied: nebulizer on, 2.5 bar; dry gas, nitrogen, 11 L min-1, dry temperature 220°C; capillary voltage, 4500 V; end plate offset, 500 V; funnel 1 radio frequency (RF), 200 Volts peak-to-peak (Vpp); funnel 2 RF, 220 Vpp; in-source collision-induced dissociation (CID) energy, 0.0 eV; hexapole RF, 120 Vpp; quadrupole ion energy, 4 eV; quadrupole low mass, 100m/z; collision gas, nitrogen; collision energy, 10 eV; prepulse storage, 7 μs. Stepping: on; basic mode; collision cell RF, from 400 Vpp to 1000 Vpp; transfer time, from 30μs to 70μs, timing; 50 %/50 %, collision energy for MSMS, 80 %, timing 50%/50%. Data dependent CID settings: intensity threshold 600, cycle time, 1 sec, active exclusion on after 2 spectra, release after 0.5 min, smart exclusion, off, isolation and fragmentation settings, size and charge dependent, width 3-15 m/z, collision energy 20-30 eV, charge states included: 1z, 2z, 3z.

Calibration of the m/z scale was performed for individual raw data files on sodium formate cluster ions obtained by automatic infusion of 1.66 μL/min of 10 mM sodium format solution of NaOH in 50/50 (v/v) isopropanol water containing 0.2% formic acid at the end of the gradient (HPC mode).

Three mixed QCs were prepared (QC roots, QC leaves, QC exudates) and run after each 8th sample. A mix of 8 commercial standards was also run after 8 samples (MM8, (Böttcher et al., 2007)). We used the following blanks: Injection blanks (ACN); extraction blanks (empty reaction tube, for roots and leaves; for exudates; empty reaction tubes with 1ml water and 1 ml methanol), furthermore for exudates: water and collection buffer from the field campaign stored in the polypropylene cryo vials as well as empty cryo-vials, all passed through the whole shipment, freeze-drying and exudate preparation process. Roots, leaves and exudates were prepared and measured in separate batches to avoid cross-contamination. Within the tissues, all samples were randomized throughout the complete process of processing and measurement.

### Data processing

The LC-qToF-MS data were processed all together with Bruker Compass MetaboScape Mass Spectrometry Software, Version 5.0.0 (Build 683) (Bruker Daltonik GmbH, Bremen, Germany). Mass recalibration, peak picking, peak alignment, region complete feature extraction, and grouping of isotopes, adduct and charge states was performed with the T-ReX algorithm in Metaboscape. Settings: Peak detection: intensity threshold, 1000 counts, minimum peak length, 7 spectra, feature signal, intensity. Minimum peak length for recursive feature extraction, 5 spectra. Retention time range, 0-18 min. Mass range, 90-1600 m/z. MSMS import method, average, grouped by collision energy. Ion deconvolution: EIC correlation, 0.8, primary ion, [M+H]+, seed ions, [M+Na]+, [M+K]+, [M+NH4]+, common ions, [M+H-H2O]+. T-ReX Positive Recalibration Auto-Detect. Feature filters: Minimum number of samples: present in 3 of 302, minimum for recursive feature extraction: present in 3 of 302, group filter: present in at least 80% of at least one group (1 group = all replicates from 1 diversity level within each species and tissue). Features from Blanks were excluded when the maximum signal in samples divided by maximum signal in blanks was ≤ 3. Quality checks included stability of retention time and signal intensity, check for carry-over, check for correct species identity. With these settings two feature tables were created: One that classified the plant diversity around the target trees by the diversity of the experimental plot (“plot diversity) and one that classified it by the local neighbourhood (“local diversity”). After processing, the data set for plot diversity contained 39077 features in total, to which 22520 fragment spectra were assigned; 3883 features from blanks were removed, leaving 35194. For local diversity: total, 38105, assigned fragment spectra 22104; 3649 features from blanks were excluded, leaving 34456.

### Statistical Analysis

For further statistics, the feature tables were exported to Metaboanalyst 4.0 (Chong et al., 2019). For the general comparison by principal component analysis (PCA) of the species-specific metabolomes (Fig 1, PCA) all species and tissues were analysed together. The data were IQR filtered before and Pareto scaled. Chow-Ruskey diagrams were produced with “intervene” (https://asntech.shinyapps.io/intervene/; (Khan and Mathelier, 2017) counting a feature as “present” when it was detected in at least 2 samples with an intensity of >/= 1000.

**Figure 1.**
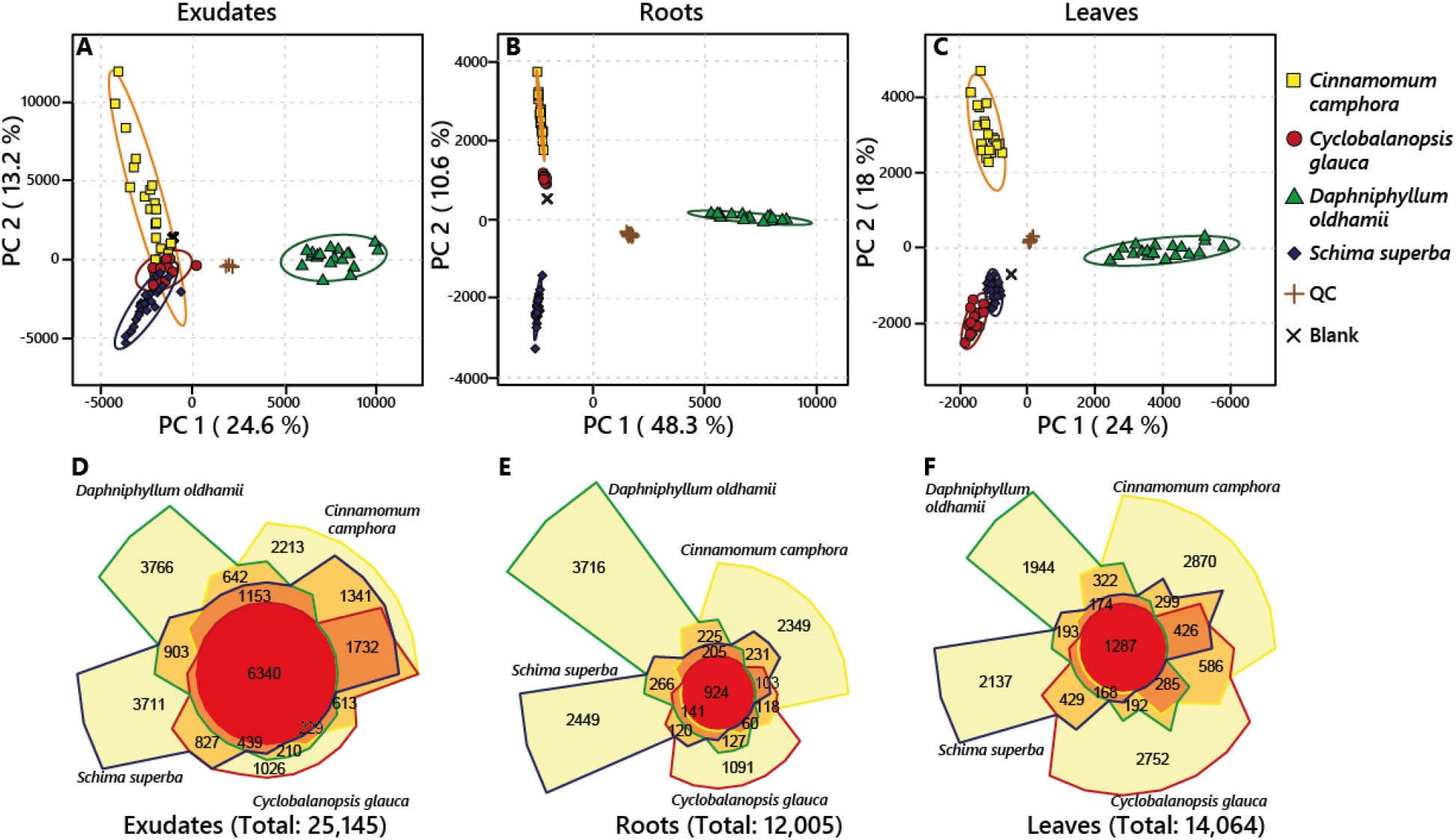
Principal Component Analysis (PCA) score plot of the first two principal components (PC) of features found in the exudates (A), roots (B) and leaves (C) of *Cinnamomum camphora* (yellow squares)*, Cyclobalanopsis glauca* (red circles)*, Daphniphyllum oldhamii* (green triangles) and *Schima superba* (blue diamonds). Quality control samples (QC, brown crosses) cluster together, demonstrating the reproducibility of the chromatographic separation; so do blank samples (black crosses). Numbers in brackets are the percentage of variation explained by the corresponding PC. Ellipses are the 95% confidence interval. Data was Pareto scaled. Second row: Venn diagrams (Chow-Rusky type) of species specific and shared features in the exudates (D), roots (E) and leaves (F) of the four different tree species showing the number of mass features unique to a single species (light yellow) or shared between two (light orange), three (dark orange) or all species (red). The area of single sections corresponds to the proportion of the number of features compared to the total number.

For Partial least squares discriminant analysis (PLS-DA) we created smaller feature tables including only the respective species and tissue metabolomes to avoid data sets with too many zeros. The date was also IQR filtered before and Pareto scaled. To group the samples in different diversity levels for PLS-DA, we used either the theoretical plot diversity or the actual local diversity. Because local diversity is maximized to 11 (See also Table S1) and varied due to tree dieback or misplantings, some diversity levels had 2 or fewer replicates. In order to have at least n= 3 per local diversity level, we grouped the samples as follows: 1 species (monoculture), 2 species, 3 or 4 species, and 5 to 7 species mixtures. In the case of *Cy. glauca*, the latter two levels (3,4 and 5) had to be combined to obtain sufficient number of replicates, and there were no trees with local diversity level of 6 or 7. For *D. oldhamii*, the monoculture and 2 species samples were combined, because one of the monocultures was invaded by another tree species, leaving only 2 replicates for monocultures. To compare the data matrices, we performed multiple response permutation procedure (MRPP) with the r-package *vegan* (v.2.5-6; https://CRAN.R-project.org/package=vegan) in R (version 3.6.0, (RCoreTeam, 2020)).

We calculated metabolite richness as the sum of all features with an intensity >1000 and metabolite diversity as the Shannon diversity of all features with an intensity >1000 using the r-package *vegan*. Two-way ANOVA, with species and plot or local richness as fixed factors, and Spearman’s rank-order correlation of metabolite richness or Shannon index and diversity were performed with SigmaPlot 14.0 (Systat Software Inc., San Jose, CA, USA).

### Compound annotation

First, all fragment spectra were matched in a parallel search against the following data bases: NIST17 (The NIST Mass Spectrometry Data Center, U.S. Department of Commerce, Gaithersburg, MD, USA), MoNa (https://mona.fiehnlab.ucdavis.edu/), ReSpect (Sawada et al., 2012), Riken public databases http://prime.psc.riken.jp/compms/msdial/main.html#MSP), Mass Bank EU (https://massbank.eu/MassBank/Index), GNPS (https://gnps.ucsd.edu/) and an internal data base (Döll, unpublished), using the MetaboScape Spectral Library Search function (Parameters: Filter: exact match of data base entry to precursor mass. Tolerances (narrow-wide): m/z 10-30mDa, mSigma 20-100, MS/MS score 900-800) Thereafter, we performed a manual annotation of species-specific compounds with features from literature and KNApSAcK database (see “Target tree species”) based on sum formula generation (Smart Formula algorithm, MetaboScape), in-source fragmentation patterns and spectral similarities to the mentioned databases. From these manual annotations, a small spectral library was created (197 species specific compounds). This library was matched against the whole data set with the MetaboScape Spectral Library Search function without the exact match of the precursor mass to find more species specific compounds similar to the already annotated ones. For the classification approach, the whole data set was searched against NIST17 without the exact match of the precursor mass. Tolerances were as above, limitation to NIST 17 was done because of computational limitations. 5. Relevant features selected after PCA (features with the highest loading on the separating principal components) or PLS-DA (20 highest VIPs – variable importance in projection on the first component) were manually annotated as above.

Annotated compounds were classified using an R workflow (Supporting Information, R Script). In brief, the annotated feature table was exported in .csv format including CAS numbers, if available, and read into R (v 4.0.3, (RCoreTeam, 2020). We employed “webchem” (Szöcs et al., 2020) packages for automatic collection of the PubChem Compound Identification number (CID). In a subsequent step, the CID was then used to retrieve descriptors (SMILES and InChIKeys) from the PubChem database. The actual classification was performed employing the ClassyFire tool (Feunang et al., 2016) using the “RAMClustR” package (Broeckling et al., 2019) in R for getting the chemical ontology information of the annotated compounds. Thus, 8101 features were classified into “Kingdom” (e.g. “Organic compounds”), “Superclass” (e.g. “Phenylpropanoids and polyketides”) and “Class” (e.g. “Coumarins and derivatives”). Sunburst plots were created in Excel 2019, aided by an R script (packages "tidyverse", "fs", "readxl", "writexl"; see supplemental file: code_snippets_data analysis_MTBLS1968.R), by adding up the feature intensities for each feature from all the samples of one species and tissue.

## RESULTS

### Metabolome composition of exudates, roots and leaves is defined by species identity

Principal component analysis (PCA) showed that the metabolomes of the four tree species can be distinguished in all three tissue types sampled (Fig. 1 A-C). In general, the metabolome of exudates showed a larger variation indicated by wider confidence bands and larger score distances within the species (Fig 1A) than leaf and root metabolomes (Fig 1B-C). The total number of mass features (25,145) in root exudates was more than twice as high as in root (12,205 in Fig. 1 D-F) or shoot samples (14,064; Fig. 1 D-F). However, root exudates of the four species shared about 25% of their metabolites, whereas in roots only 7.7% and in shoots 9.1% of all the metabolites were found in all four species.

The metabolomes of *D. oldhamii* always clearly separated on the first principal component (PC) axis. The compounds responsible for separating *D. oldhamii* exudates and roots from the other species are likely the Daphniphyllum alkaloids, which had the highest loadings on PC1 (Table S2 and S3; Fig. 2B and 2C, third column). Interestingly, the leaf metabolomes of *D. oldhamii* contained relatively few alkaloids (Fig 2D, third column), the discriminating compounds were glycosides, most likely with a flavonoid backbone, a disaccharide, a putative coumarin and a steroidal compound (Table S4). The other three species mainly separated on PC2 (Fig 1 A-C). Whereas the exudate metabolomes (Fig. 1A:) of *C. camphora*, *Cy. glauca* and *S. superba* showed some overlap, their leaf and root metabolomes clearly separated (Figs 1 B-C;).

**Figure 2.**
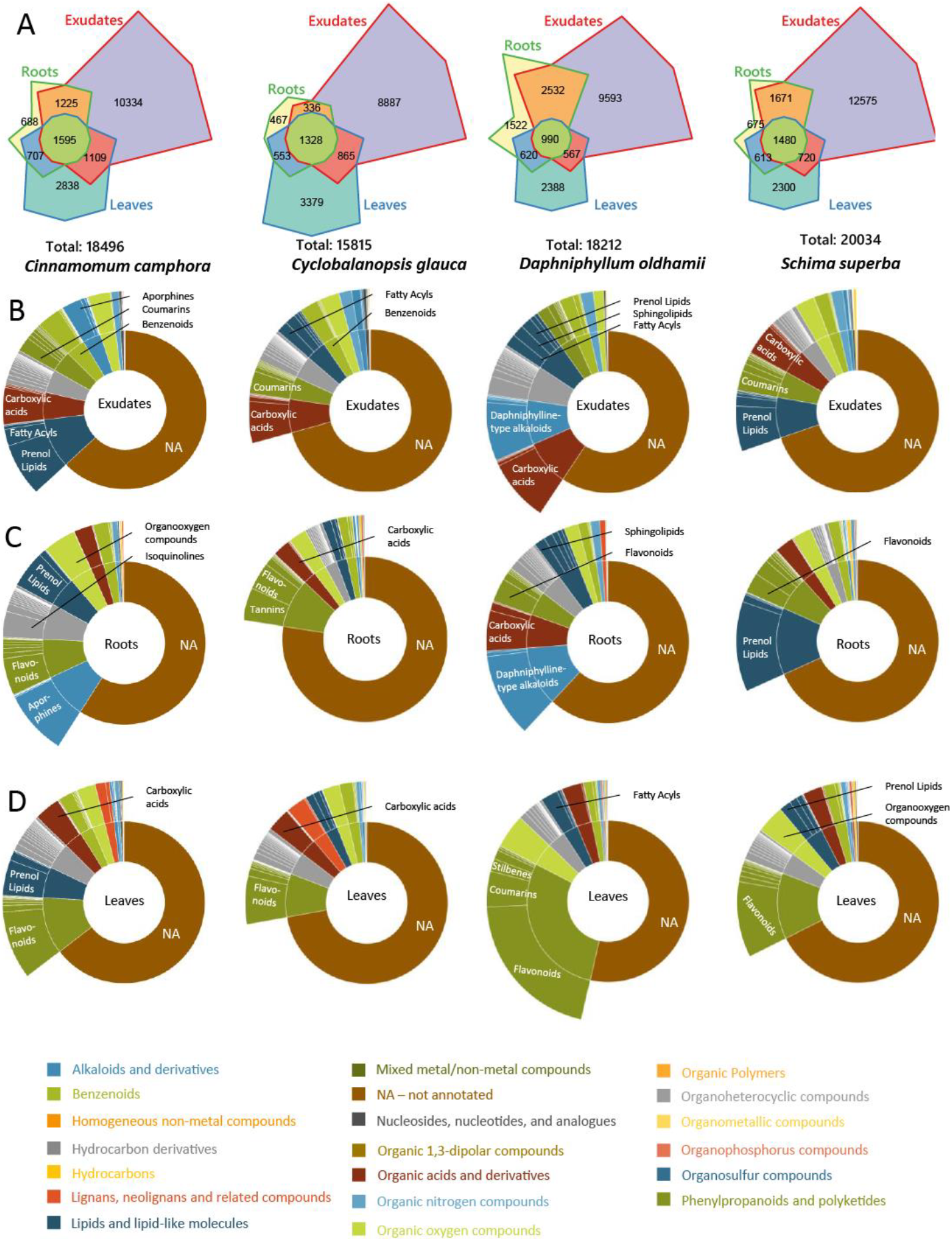
Chow-Rusky Venn diagrams (A) of features in the metabolomes of the four tree species *Cinnamomum camphora, Cyclobalanopsis glauca, Daphniphyllum oldhamii* and *Schima superba* that are unique to exudates (red), roots (green) and leaves (blue) and features that are shared between two or all of the different samples. The area of single sections corresponds to the proportion of the number of features compared to the total number. Second to fourth row: Sunburst Plots visualizing the metabolome compositions of exudates (B), roots (C) and leaves (D). The inner circle represents the *chemical superclass* according to the *ClassyFire* hierarchical classification while the outer circle represents the *chemical class*. Proportions of the circle represent the sum of the intensities of all features assigned to *chemical class*. *Chemical superclasses* are defined in the legend at the bottom of the figure and selected *chemical classes* are specified in the sunburst plots.

The presence of laurolitsine-like alkaloids separated both *C. camphora* exudates and root metabolomes from the other species (Table S2 and S3). Several terpenoids and a putative phenylpropanoid in *C. camphora* exudates (high loadings on PC2; Supplemental Table S2); and a procyanidin, a putative reticuline, a sesquiterpene and an unknown metabolite in *C. camphora* roots were the most responsible for separating this species from the other two. The *C. camphora* leaves were typified by specific terpenoids. In *S. superba* roots and root exudates, saponines had the highest influence in separating *S. superba* from the rest (Table S2 and S3). A coumarin (in exudates) and a putative phenylpropanoid (roots) contributed to the separation. For *Cy. glauca* leaves, quinic acid, polyphenols and terpenoids were the metabolites with the highest loading and therefore discriminating these metabolomes from the other species. We could not identify discriminating features for roots and root exudates of *Cy. glauca*, because there was too much overlap with the metabolomic profiles of *C. cinnamonum* and *D. oldhamii* in PC1 vs. PC2 (Fig 1A).

When comparing samples within species, it became apparent that in all four species the exudates had the highest number of unique mass features, followed by leaves and roots (Fig. 2A). The number of root specific features was generally less than 4%, with the exception of *D. oldhamii* (8.4%). In *D. oldhamii, C. camphora* and *S. superba,* the numbers of features shared between roots and exudates were greater than those shared between roots and leaves (Fig. 2A)

In other to better characterize the metabolomes of exudates, roots, and leaves per species we used an automated approach to classify our tentatively annotated features in a chemical class by using the ClassyFire tool ((Feunang et al., 2016); see methods section). This tool uses structural information (e.g. SMILES, InChIs or IUPAC names) to categorize chemical entities into hierarchical chemical classes. Thus, we were able to categorize between 22% (roots *Cy. glauca*) and 46% (leaves *D. oldhamii*) of the m/z features into main compound classes (Table S7). The resulting plots reveal that carboxylic acids are a major common compound class in root exudates (Fig 2 B). In roots, specific alkaloids, flavonoids and prenol lipids are among major compound classes we could categorize. The leaf metabolomes of all four species are dominated by phenylpropanoids with flavonoids as the most abundant chemical glass, followed by organoheterocyclic compounds (e.g. imidazopyrimidines, indoles, azaspirodecanes) or organooxygen compounds (e.g. alcohols and polyols, carbohydrates and conjugates, carbonyl compounds) (Fig. 2 D).

Prenol lipids were omnipresent, but most prominently present in roots and exudates of *S. superba* as well as in exudates and leaves of *C. camphora* (Fig. 2 B-C). Aporphines, a class of alkaloids, and the structurally related isoquinolines, were most prominent in *C. camphora* roots, but less in the exudates, whereas the classified metabolome of both *D. oldhamii* roots and root exudates consisted for a large part of the typical Daphnipyllum alkaloids (Fig. 2 B-C, 3^rd^ column).

Interestingly, neither in *C. camphora* nor in *D. oldhamii* leaves, alkaloids were as pronouncedly present as they were in their exudates or roots. The plots also show that on the level of compound class, the chemical profiles of roots and root exudates are more similar to each other than to leaf metabolomes (Fig. 2B-D). This is also in line with the high number of overlapping features for roots and exudates (Fig. 2A). Finally, we found coumarins to be present in *C. camphora* and *S. superba* exudates, as well as in *D. oldhamii* leaf metabolomes. Sphingolipids formed a notable part of *D. oldhamii* roots and root exudates, whereas the roots of *Cy. glauca* showed tannins and flavonoids as major classes (Fig. 2D).

### Effects of plot and local tree diversity on metabolome composition

Because the metabolomes differed substantially among species and tissues types, we assessed the effect of plot and local tree diversity by species and sample. Partial least squares discriminant analysis (PLS-DA) showed that plot diversity affected the metabolome composition dependent on the sample type as well as on species identity. The exudate metabolomes of three of the four tree species (*Cy. glauca, D. oldhamii* and *S. superba*) were significantly affected by plot diversity level (Fig. 3; MRPP P < 0.05; Table S5).

**Figure 3.**
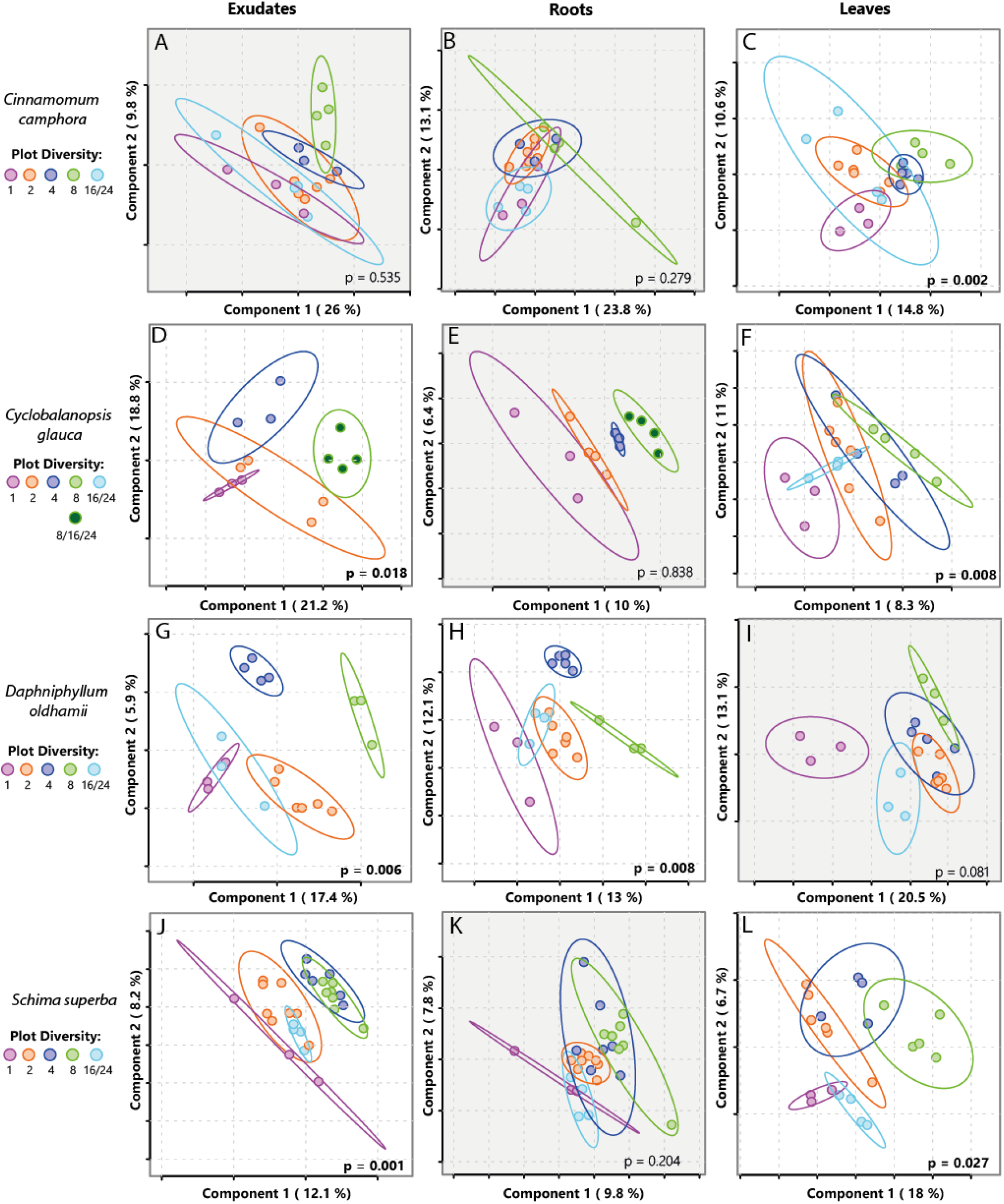
Partial Least Squares – Discriminant Analysis (PLS-DA) score plots of the features found in the exudates, roots and leaves of the 4 tree species *Cinnamomum camphora* (A-C)*, Cyclobalanopsis glauca* (D-F)*, Daphniphyllum oldhamii* (G-I) and *Schima superba* (J-L). Different colors indicate the tree species diversity in the investigated plot. Percentages at the axes indicate the variation explained by the single components. Ellipses are the 95% confidence interval. P – values are global values and are based on pairwise multiple-response permutation procedures (MRPP). Because of an insufficient number of replicates, diversity groups 8 and 16/24 in *Cyclobalanopsis glauca* were combined for statistical analysis. The data was Pareto scaled and features from blanks removed. P – values smaller than 0.05 are in bold and the corresponding plot is highlighted.

In the roots, only the metabolomes of *D. oldhamii* varied significantly with plot diversity level, whereas the leaf metabolomes of *C. camphora*, *Cy. glauca* and *S. superba* all varied significantly due to plot diversity The effect of local diversity was less pronounced; only *S. superba* exudates showed a significant difference (Fig. 4), while none of the root metabolomes was affected significantly. The leaf metabolomes of the same three species (*C. camphora, Cy. glauca* and *S. superba*) responding to plot diversity, also showed a significant response to local diversity (Fig. 4). Neither for plot nor for local diversity, any of the pairwise comparisons in the MRPP analyses showed statistically significant differences (Table S5 and S6).

**Figure 4.**
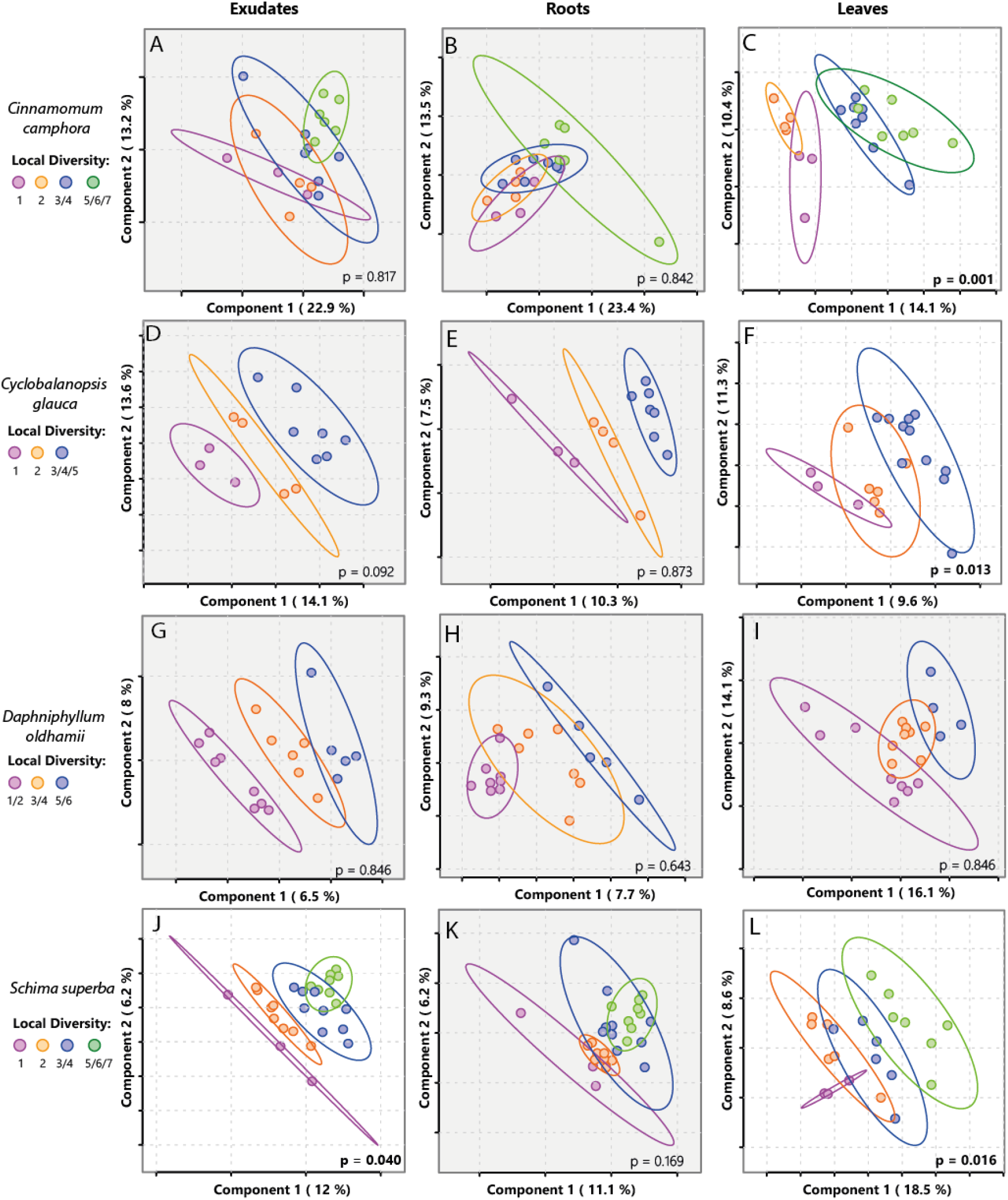
Partial Least Squares – Discriminant Analysis (PLS-DA) score plot of the first two components of the features found in the exudates, roots and leaves of the 4 tree species *Cinnamomum camphora* (A-C)*, Cyclobalanopsis glauca* (D-F)*, Daphniphyllum oldhamii* (G-I) and *Schima superba* (J-L). Different colors indicate the actual tree species diversity in the immediate local neighbourhood of the target tree (as opposed to “plot diversity”). Percentages at the axes indicate the variation explained by the single components. Ellipses are the 95% confidence interval. Because of an insufficient number of replicates, some of the diversity groups in “Local Diversity” were combined for statistical analysis. P – values are global values and are based on pairwise multiple-response permutation procedures (MRPP). The data was Pareto scaled and features from blanks removed. P – values smaller than 0.05 are in bold and the corresponding plot is highlighted.

For all species and samples that showed a significant overall response to plot or local diversity, we picked the features with the highest VIP (variable importance in projection) value to analyse their response to tree diversity in more detail (Table S8, S9, and S10). Several of these features were annotated to compound classes that also constituted a large fraction of the respective metabolomes. In the exudates and roots of *D. oldhamiii,* two of the typical Daphniphyllum alkaloids were responding to plot diversity, and in the exudates of *S. superba* the levels of saponins and phenylpropanoids varied with plot and local diversity (Fig. 5 B and D). In the leaves of *C. camphora*, *Cy. glauca* and *S. superba,* flavonoids (e.g. kaempferols, rutin and other quercetins) and terpenes were among the compounds responding to variation in diversity level.

**Figure 5.**
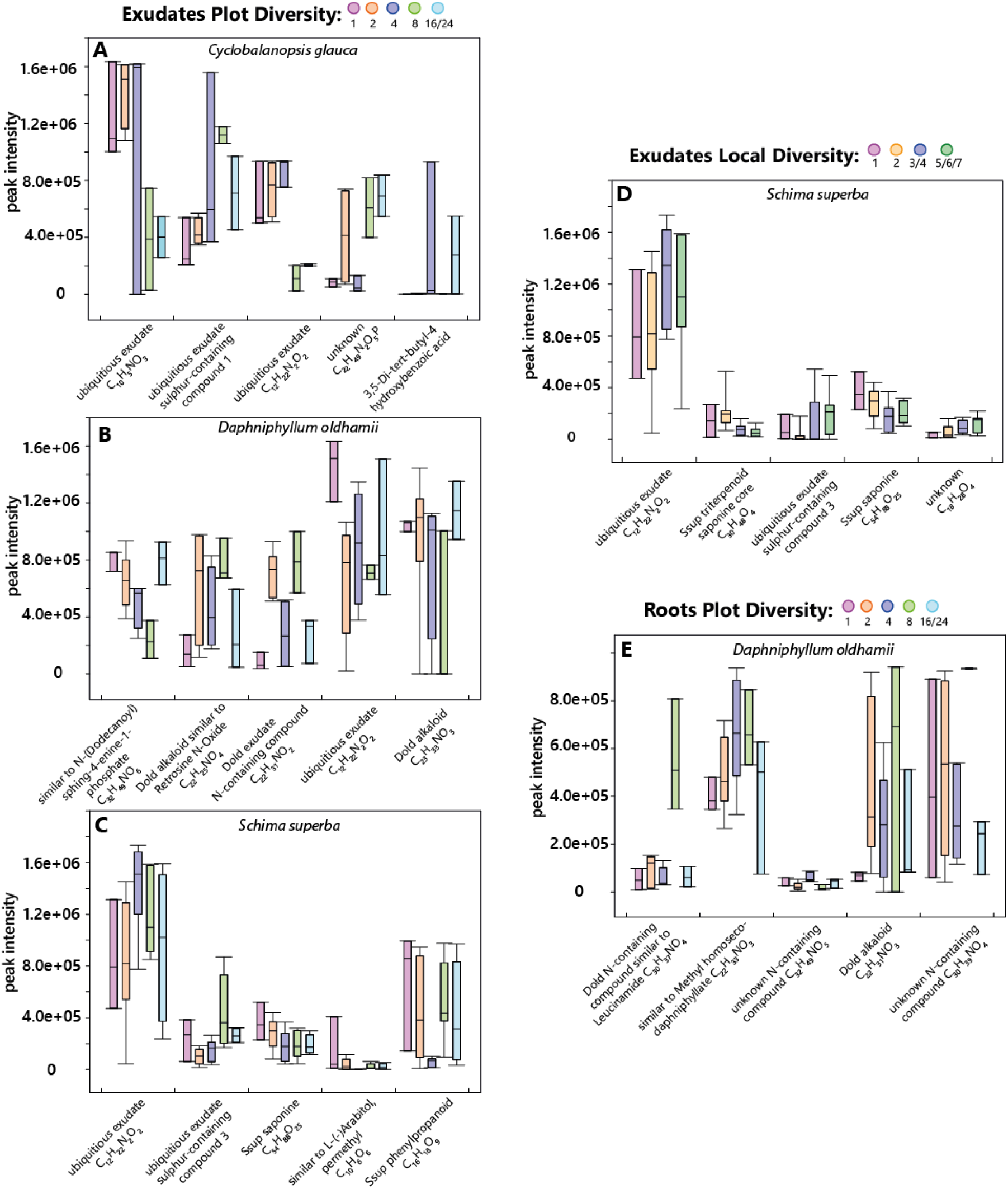
Boxplots of the top 5 VIPs (Variable Importance in Projection) of PLS-DA that showed a significant global difference in the metabolome (MRPP P < 0.05) in Fig. 4 (“Box”: 25^th^ to 75^th^ percentile and median, “whiskers”: 10^th^ and 90^th^ percentile).

Despite the presence of features responding to biodiversity level, there was no overall uniform response on species, sample or compound level. For example, in *S. superba* leaves, quinic acid decreased in intensity with plot and local diversity, while in the same species rutin was increasing with increasing local or plot diversity (Fig 6 C and F). Daphniphyllum alkaloids either showed a hump-shaped response or rather decreased at intermediate plot diversity levels in *D. oldhamii* exudates (Fig 5 B). In roots, they either peaked at diversity level 8, or increased with diversity (Fig 5 E). Within species and samples, we identified some common patterns. Saponin levels in *S. superba* commonly decreased (Fig. 5 C and D), whereas a farnesene-like sesquiterpene in *C. camphora* consistently increased with local and plot diversity (Fig. 6 A and D). In general, the peak intensity of the single features varied substantially across species and sample types, likely due to sampling issues and low replication rates.

**Figure 6.**
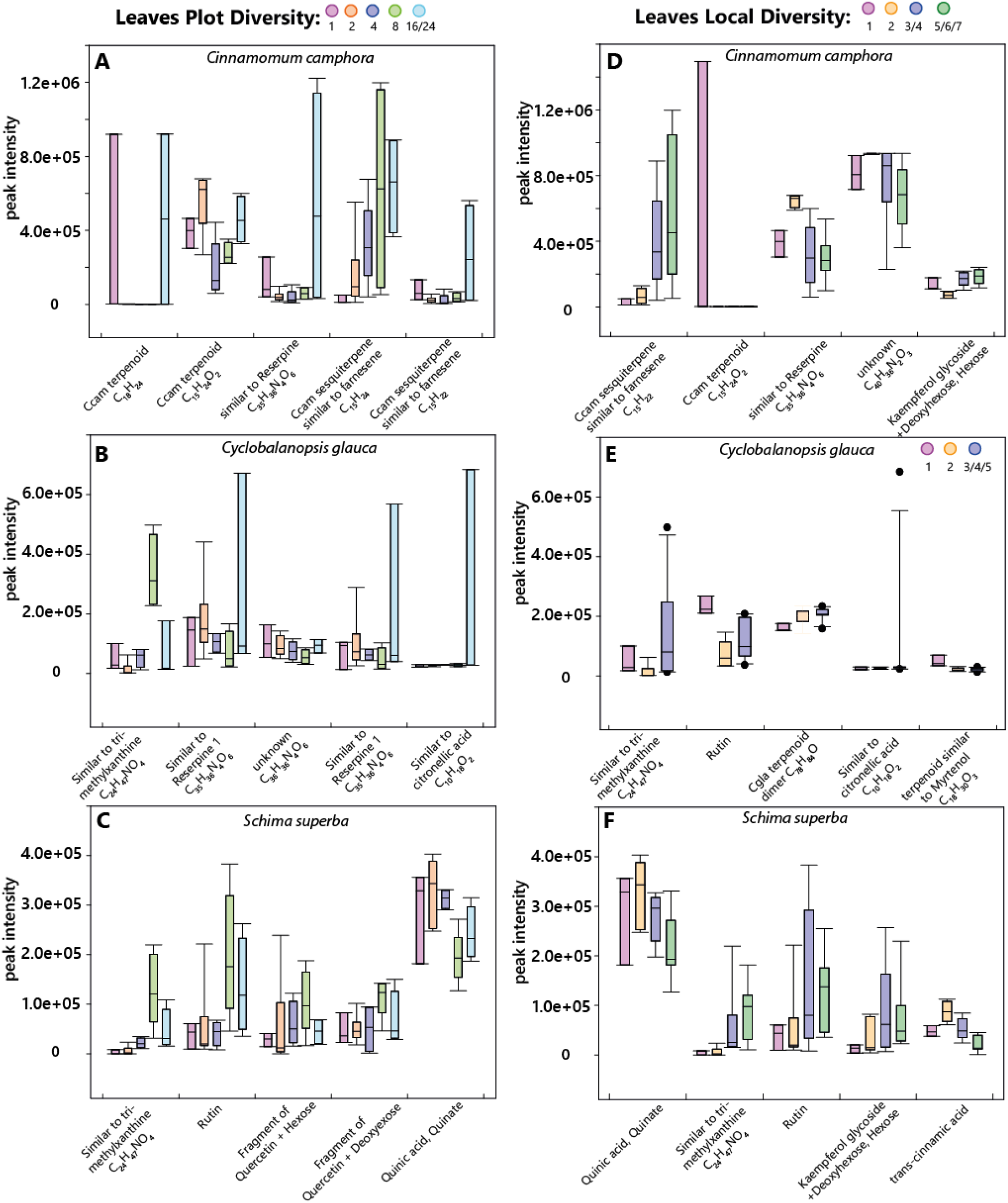
Boxplots of the top 5 VIPs (Variable Importance in Projection) of PLS-DA that showed a significant global difference in the metabolome (MRPP pP< 0.05) in Fig. 4. (“Box”: 25^th^ to 75^th^ percentile and median, “whiskers”: 10^th^ and 90^th^ percentile, circles: outliers)

### Effects of plant diversity on species chemical diversity

Finally, we correlated chemical richness, *i.e.* the number of mass features with intensity > 1000, or the chemical diversity, calculated as the Shannon diversity of the mass features, with either local diversity or plot diversity. Spearman rank-correlations revealed that metabolite richness was not significantly affected by plot diversity in any sampling type or tree species. The leaf metabolite richness of *Cy. glauca* (P = 0.053) and *D. oldhamii* (P= 0.071) showed a marginally significant response, but in opposite directions. Metabolite richness increased with plot diversity in *Cy. glauca* leaves, and decreased with plot diversity in *D. oldhamii* (Fig. S3; third column). Local diversity only affected leaf metabolomes of *Cy. glauca* (P= 0.049) and marginally so, the exudates of *S. superba* (P= 0.056). In both cases, local diversity positively correlated with metabolite richness (Fig. S4). In none of the samples we found a significant correlation between Shannon diversity of mass features and local or plot diversity level (Figs S5 and S6).

## DISCUSSION

Overall, exudates, roots and leaves each showed clearly species-specific metabolomic profiles. Leaf and root metabolomes were more distinctive than root exudates. The prominent presence of specific alkaloids separated the *C. camphora* and *D. oldhamii* root and root exudate metabolomes from each other as well as from the other two tree species. As postulated, root and root exudate metabolomes were more similar to each other than to leaf metabolomes. Carboxylic acids formed a large part of all exudates, whereas phenylpropanoids, in particularly flavonoids, dominated the classifiable subset of all leaf metabolites. All four tree species showed a metabolomic response to tree diversity in at least one of the metabolomes sampled. Both root exudates and leaf metabolomes responded more often to differences in tree diversity levels than roots. Plot diversity level had an overall larger effect on tree metabolomes than local diversity. Important features driving the differences in the metabolomes were saponines in *S. superba* exudates *, D. oldhamii* specific alkaloids in roots, kaempferols, quercetins and quinic acid in *S. superba* leaves, and sesquiterpenes in *C. camphora.*

Previous studies have shown that leaf metabolomes of various tropical species show a high level of species-specificity (Richards et al., 2015, Salazar et al., 2018, Sedio et al., 2019, Sedio et al., 2017). The metabolomic diversity among congeners can be greater than chemical diversity among genera (Sedio et al., 2019). Here we showed that also roots and their exudates have distinct species-specific metabolomic profiles. Compared to leaves and roots, however, exudates had higher numbers of specific features with a higher intra-specific variance. Moreover, the profiles showed more overlap, despite the fact that we normalized peak areas over NPOC to mitigate differences in exudation rates. Likely this is due to the fact that exudates were sampled *in situ,* which means that these samples inevitably contained compounds from microbes and organic matter in the rhizosphere. Additionally (local) differences in nutrient status among plots unrelated to plot diversity may have caused differences in exudation patterns (Meier 2020). Commonly, much less than 10% of the mass features in metabolomic experiments can be assigned, even in model systems (da Silva et al., 2015). This limitation prohibits the ecological interpretation of differences in metabolomic profiles (Peters et al., 2018). By using bioinformatics tools to compare m/z features to online metabolite databases, we could broadly classify up to 45% of the features to compound classes (Figure 2). This did not only allow us to derive broad ecological functions, but also created a starting point for more targeted analyses combined with biological experiments to assess the ecological relevance of particular metabolites in more detail.

Carboxylic acids were one of the most abundant metabolite classes in the root exudates (van Dam and Bouwmeester, 2016). Other carboxylic acids (mono-, di-, and tricarboxylic acids) are found in the root exudates of birch and spruce (Sandnes et al., 2005). It is known that carboxylic acids, like citrate or malate, play a role in P mobilization (Inderjit and Weston, 2003). In *D. oldhamii* the second largest (in exudates) and largest group (in roots) of annotated features were alkaloids. Daphniphyllum alkaloids are structurally diverse group of metabolites, which were isolated from the bark of *Daphniphyllum spp*. and were also reported in leaves and fruits (Wu et al., 2013). Therefore, it is surprising that we found Daphniphyllum alkaloids in roots and root exudates, but were not or only scarcely present in leaves. We identified one study showing that these alkaloids may have insecticidal activities (Li et al., 2009). Because Daphniphyllum alkaloids are mostly studies for their potential anticancer properties, little is known about their ecological roles. The exudates and roots of *C. camphora* contained a considerable number of lauralitsine and reticuline-like alkaloids. These alkaloids, which are also mainly studied for their medicinal properties, were previously reported to be present in the roots (Custódio and Florêncio da Veiga Junior, 2014), but not in the leaves. Their ecological functions are not experimentally assessed. Considering that alkaloids in general serve as anti-herbivore defences (Mithofer and Boland, 2012), it is likely that the alkaloids in *D. oldhamii* and *C. camphora* roots and exudates serve as defences against root feeders and pathogens. Further experiments, preferably under controlled conditions, are needed to falsify this hypothesis. Prenol lipids made up a substantial fraction of annotated features in the exudates, roots and leaves of *C. camphora* and *S. superba.* This class contains molecules consisting of one or several isoprene (C5) units (Fahy et al., 2005). It includes mono (C10) – and sequiterpenoids (C15), which are known for their roles in direct and indirect defence against herbivores and pathogens. In addition, also carotene (C40) which plays a role in photosynthesis (Fahy et al., 2005) and saponines which have been shown to modulate soil microbial communities (Fujimatsu et al., 2020), belong to this group. The roots of *Cy. glauca* contained tannins as a major class. Tannins are typical for the oak family, including *Cy. glauca* (Wakamatsu et al., 2020), which are known as defences to a broad range of herbivores (Barbehenn and Constabel, 2011). As mentioned earlier, our metabolomic analyses of these four chemically poorly described tree species is a starting point for testing their chemical ecology in more detail.

Root exudate and root metabolomes showed a high overlap in the number of features detected as well as in the chemical classes that could be annotated (Fig 2). In particular, *D. oldhamii* and *S. superba* exhibited a high similarity in root exudate and root metabolites. On the other hand, *C. camphora* shows a high overlap in the number of features, but the composition of root exudates and roots are very different. This might be due to the different ways in which metabolites are exuded by plants (Oburger and Jones, 2018) or the interspecific interactions with trees in the local neighbourhood (Xia et al., 2016). The metabolomes of root exudates shared also a number of features with the leaf metabolomes. This was not reflected in the composition of the annotated metabolome. The shared features might belong to the group that could not be annotated and therefore makes an interpretation more difficult. Leaf metabolites could be introduced into the soil via leachate from leaf litter and show up in our exudate samples, despite cleaning and washing of roots prior to sampling.

In addition to the species-specific metabolomic profiles, we also found that the metabolomes of the different species and organs responded to differences in tree diversity levels. Especially exudate and leaf metabolomes varied with plot diversity level, whereas the fine roots we sampled had rather constant profiles. The latter is in line with a recent study, showing that the glucosinolate profiles of fine roots of *Brassica* spp. did not change in response to local or systemic herbivory (Tsunoda et al., 2018). Fine roots of trees have high turnover rates, and therefore trees may invest less in defending their fine roots (Bouma et al., 2001, Yanai and Eissenstat, 2002). This is supported by the fact that we found fewer organ-specific metabolites in roots than in leaves.

Other than expected, plot diversity had an overall stronger effect on exudate and leaf metabolome profiles than local diversity. Our hypothesis was based on the fact that direct neighbouring trees would have a stronger effect on our target trees than more remote trees in the plot. The BEF China experiment was deliberately planned to have a high tree density (0.6 tree m^−2^)(Bruelheide et al., 2014), which means that ten years after planting the roots may have grown sufficiently to contact also more remote trees. Even in grasslands, where plants are growing in much closer proximity, the local neighbourhood also determined a small part of the variance in exudate composition (Dietz et al., 2019). In addition, trees may be connected via widespread mycorrhizal networks, thus influencing each other beyond the local neighbourhood (Courty et al., 2010). This may cause that overall plot diversity has a larger effect on root exudates than previously expected.

Only for *Cy. glauca* leaves we found a consistent and statistically significant increase of chemical diversity with local and plot tree diversity. Also, the number of exudate metabolites in *S. superba* showed a positive trend (P = 0.056) with local diversity, whereas the number of metabolites in *D. oldhamii* leaves tended to decrease with plot diversity. Whereas such variable relationships between diversity level and chemical richness or diversity is in line with studies analysing grassland species (Ristok et al., 2019), the high variation among samples may also have limited our ability to detect general trends Partly, this is due to the low replication level of species combinations, which is a common issue in BEF experiments manipulating plant diversity (Uthe et al., 2020). The experimental plots and as well our samples are designed to vary species diversity, whereby plots of the same diversity will differ in the composition of species (Bruelheide et al., 2014). This caused that each focal tree within diversity level was exposed to different interacting trees. This variation among plots of the same diversity level could dilute specific responses, leading to a less pronounced response in the metabolome. Additional variation can have emerged due to the vastness of the experiment and distances among (replicate) plots (Bruelheide et al., 2014). The vast spatial scale may cause differences in slope, humidity, soil characteristics et cetera, which may have affected the metabolomes. For example, substrate properties like acidity and the type of topsoil have been shown to influence the exudation in beech (Meier et al., 2020). Soil acidity is associated with nutrient availability, because a decrease in soil pH will reduce availability of N and P and also change the microbial composition in the soil (Rousk et al., 2009).

## CONCLUSION

We showed that plant diversity affects aboveground and belowground metabolomes of four subtropical tree species. Overall, plot diversity as well as the local neighbourhood has an impact on the metabolome of root exudates and leaves. Studies on tropical trees have linked metabolomic diversity to differences in insect community composition (Richards et al., 2015, Zu et al., 2020). Our observation that tree diversity affects leaf metabolomes thus may impact herbivore communities on trees of the same species, but growing in plots with different diversity levels. Similarly, differences in root exudate composition may shape the microbial diversity in the rhizosphere (Haichar et al., 2008). We also showed that the response of specific metabolites to plot diversity is dynamic and not linearly correlated to diversity gradients. This suggests that in fact not species diversity, but species composition might be the important driver of changes in plant chemical diversity. In future studies, the results of our field analyses may be used to manipulate single factors at a higher replication level to disentangle the individual contributions of species identity and neighbouring trees on tree metabolomes. These future studies may benefit from our ecometabolomics workflow to identify chemical classes of metabolites that could be investigated with more targeted approaches.

## ACKNOWLEDGMENTS

AW, ML and NMvD thank Christian Ristok for help with tree species selection, Gonzalo Garcia de Leo for doing a pre-experiment in the field, and Florian Schnabel for identifying trees. AS and SD thank Linnea Smith for her help in compiling R scripts for data base queries. All authors gratefully acknowledge funding by the Deutsche Forschungsgemeinschaft (DFG, German Research Foundation; grant 319936945/GRK2324) and the University of Chinese Academy Sciences (UCAS). AW, AS and NMvD gratefully acknowledge the support of the German Centre of integrative Biodiversity Research (iDiv) funded by the German Research Foundation (DFG–FZT 118, 202548816)

## AUTHOR CONTRIBUTIONS

AW, ML, NMvD designed the study. ML sampled exudates, root and leaf samples in the field in China, after with SD processed the samples and performed the chemical analyses. The metabolomic data were processed by AW, SD and AS. SN was responsible for data archiving and curation on MetaboLights. AW, SD, ML and NMvD took the lead in writing the paper with critical contributions of AS, XX, and SN.

## DATA AVAILABILITY

LC-MS data can be accessed at the MetaboLights data repository for metabolomics experiments (Haug et al., 2020) https://www.ebi.ac.uk/metabolights/) as MTBLS1968.

## Supporting_Information

