## Supplemental Information for "Plot diversity differentially affects the chemical composition of leaves, roots and root exudates in four subtropical tree species"

**Supplemental Figures and Captions:**

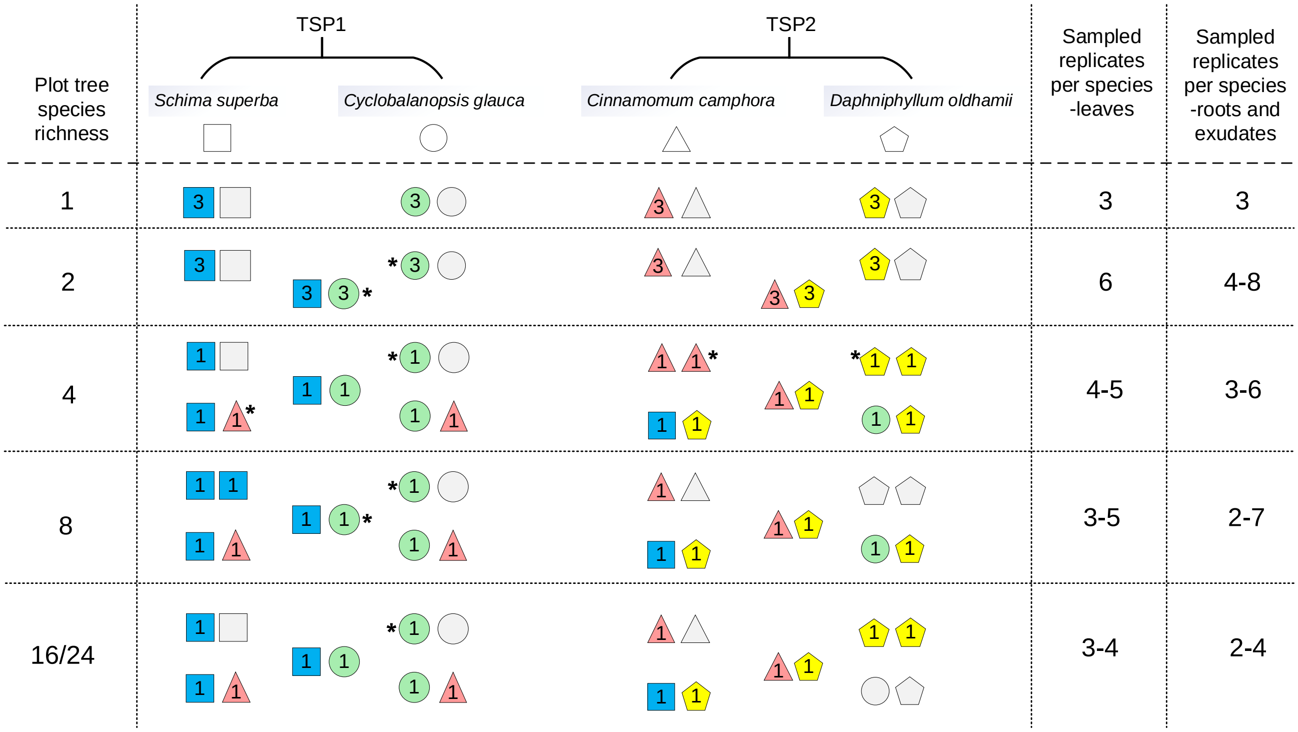

**Figure S1** Details of sampling for the 4 targeted species in the BEF-China tree diversity experiment (Site B) under different plot tree species richness levels (1, 2, 4, 8, 16/24). Different symbols indicate different species (i.e. square □ indicates *Schima superba*, circle ○ indicates *Cyclobalanopsis glauca*, triangle △ indicates *Cinnamomum camphora*, pentagon ⬠ indicates *Daphniphyllum oldhamii*). The sampled combinations of tree species pairs (TSPs) under different tree richness levels are shown. Symbols with colors mean the sampling trees, and the number inside each colored symbol indicates the replicate number of the corresponding treatment. Light grey symbols mean none sampling, as only half of the trees under monoculture were sampled and the availability of the targeted TSPs under high species richness levels were limited in the plots. Symbols with asterisk (*) indicate that one sample under the corresponding treatment was reassigned for root and exudate samples. The sampled and analyzed number of leaves, roots and exudate samples replicates (i.e. with the reassigned samples counted in) for each tree species under the five richness levels are shown in the right part of this Figure.

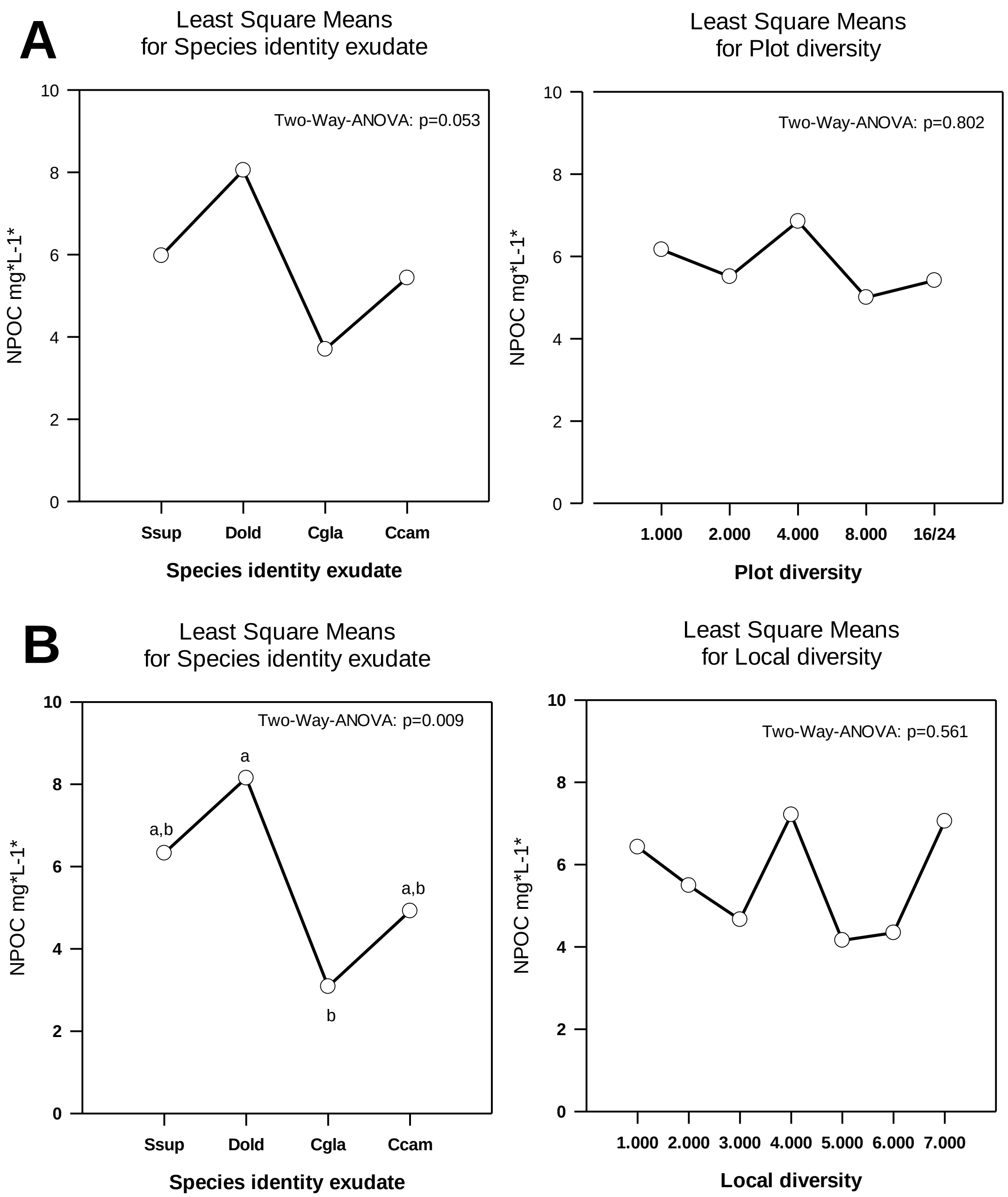

**Figure S2**: NPOC (Non-purgable organic carbon) measured in the original root exudate solutions collected from four tree species in a Chinese subtropical forest in stands with different species diversity levels. A: Main effects of Two-Way Analysis of Variance (General linear model) Factors (independent variables): Species identity (F = 2.707); plot diversity level (F=0.802). F for spec. ident. x plot diversity = 0.634. B: Main effects of Two-Way Analysis of Variance (General linear model, no interactions due to low number of replicates) with all pairwise multiple comparison procedures (Holm-Sidak). Overall significance level = 0.05 Factors (independent variables): Species identity (F = 4.153); local diversity level (F=0.816). Abbreviations: *Ssup:* Schima superba *Dold: Daphniphyllum oldhamii, Cgla: Cyclobalanopsis glauca Ccam: Cinnamomum camphora*

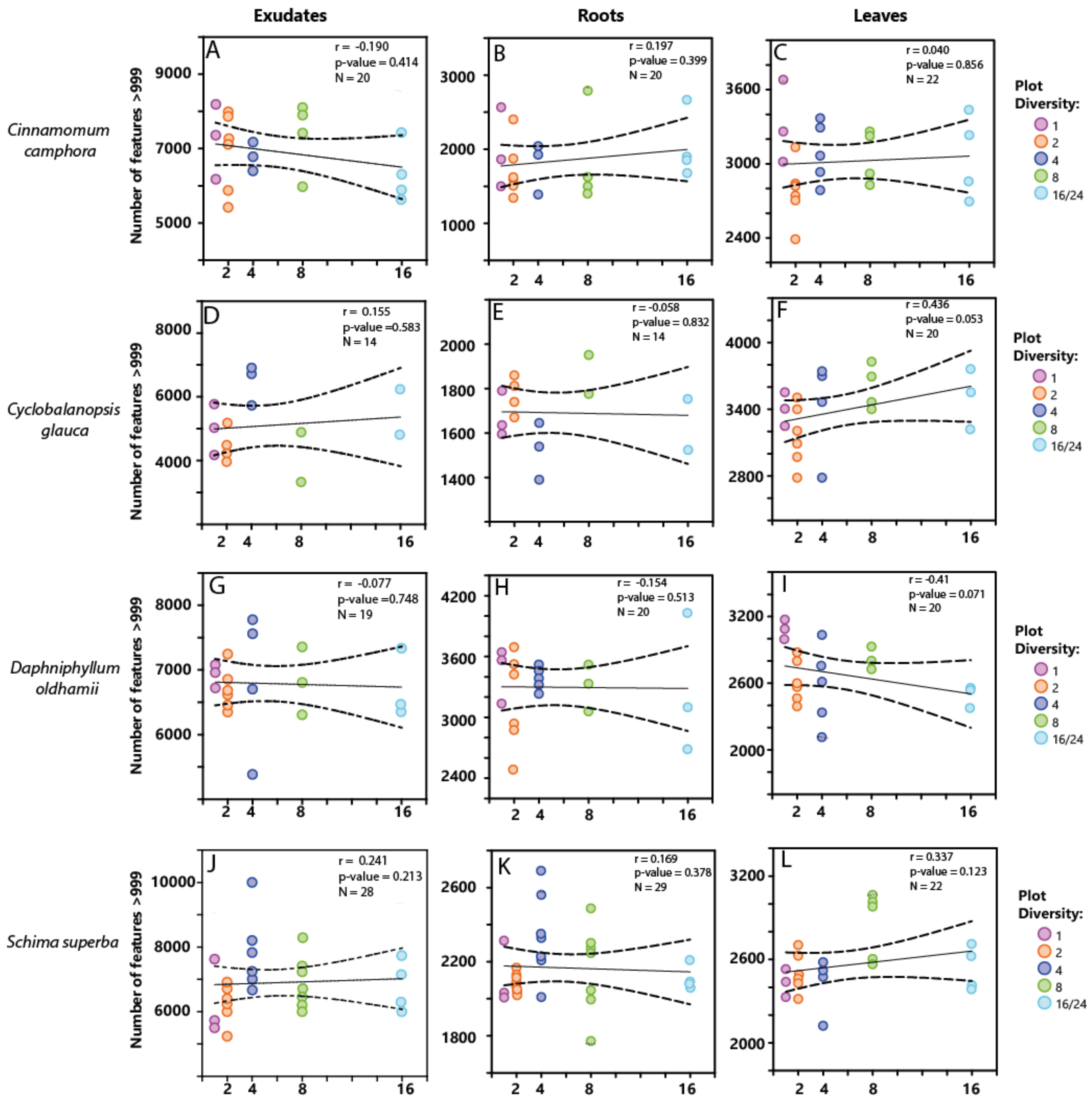

**Figure S3:** Spearman Rank Correlation of the diversity level of 4 tree species *Cinnamomum camphora* (A-C)*, Cyclobalanopsis glauca* (D-F)*, Daphniphyllum oldhamii* (G-I) and *Schima superba* (J-L) and the metabolite richness in the exudates, roots, and leaves (calculated as the sum of all features with an intensity >999). The diversity levels were either the diversity of the experimental plot. Dotted lines indicate the confidence interval of the regression (standard deviation x 3). N is the number of replicates, r is the correlation coefficient and p the p value.

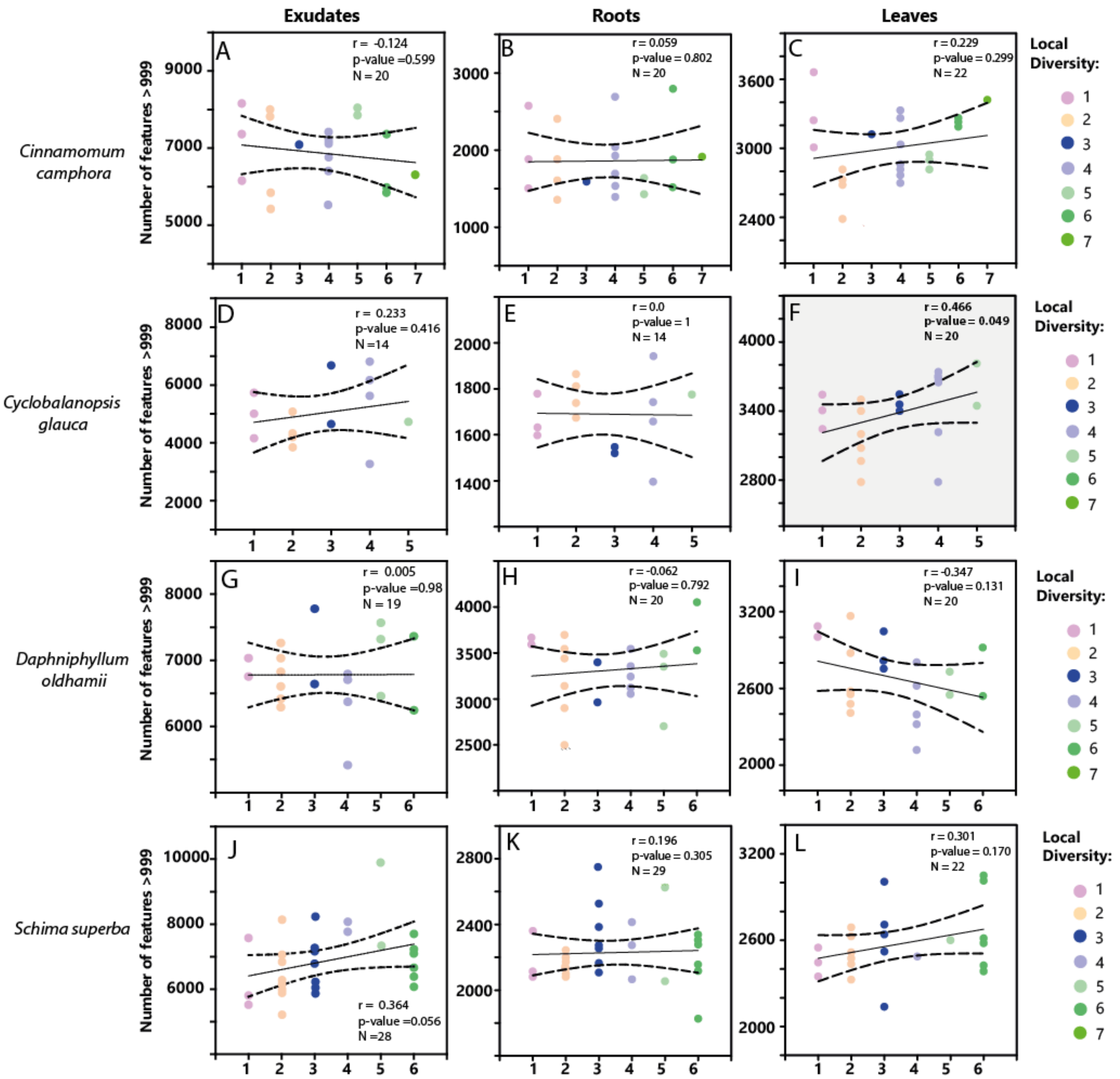

**Figure S4:** Spearman Rank Correlation of the diversity level of 4 tree species *Cinnamomum camphora* (A-C), *Cyclobalanopsis glauca* (D-F), *Daphniphyllum oldhamii* (G-I) and *Schima superba* (J-L) and the metabolite richness in the exudates, roots, and leaves (calculated as the sum of all features with an intensity >999 ). The diversity levels were the diversity level of the local neighbourhood of a single tree. Dotted lines indicate the confidence interval of the regression (standard deviation x 3). N is the number of replicates, r is the correlation coefficient and p the p value.

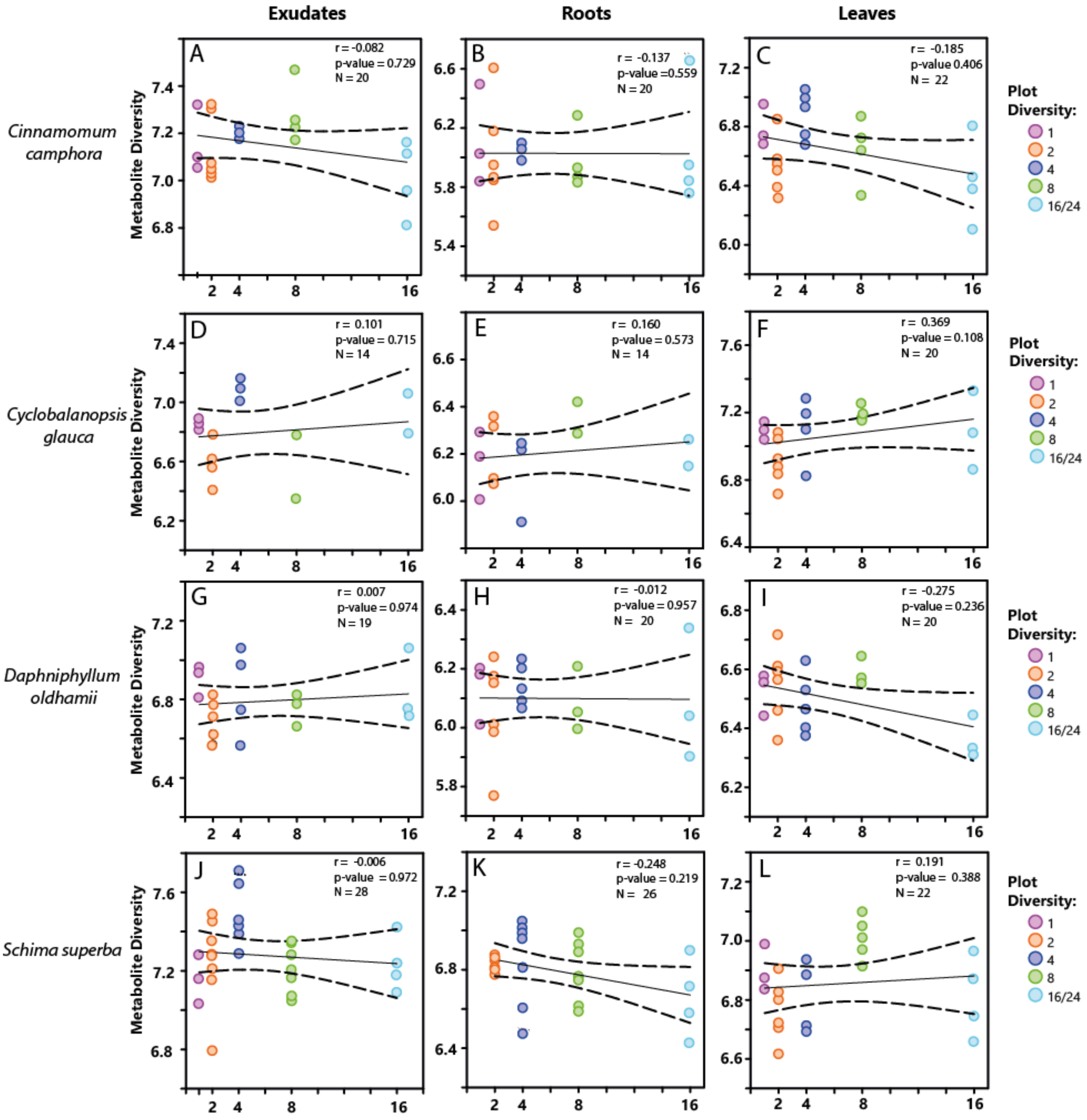

**Figure S5** Spearman Rank Correlation of the diversity level of 4 tree species *Cinnamomum camphora* (A-C)*, Cyclobalanopsis glauca* (D-F)*, Daphniphyllum oldhamii* (G-I) and *Schima superba* (J-L) and the metabolite diversity in the exudates, roots, and leaves (calculated as Shannon diversity). The diversity levels were either the diversity of the experimental plot. Dotted lines indicate the confidence interval of the regression (standard deviation x 3). N is the number of replicates, r is the correlation coefficient and p the p value.

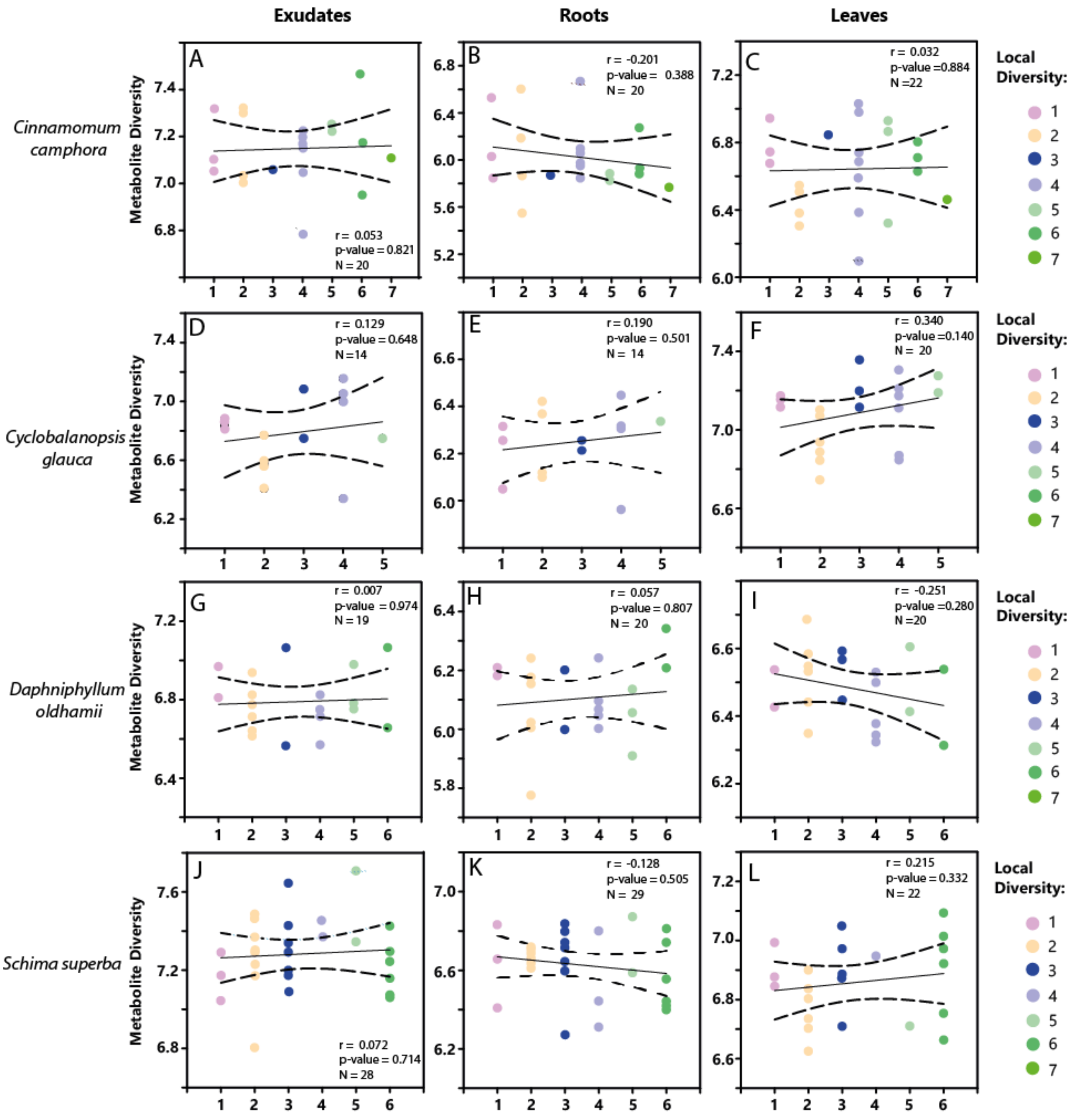

**Figure S6:** Spearman Rank Correlation of the diversity level of 4 tree species *Cinnamomum camphora* (A-C), *Cyclobalanopsis glauca* (D-F), *Daphniphyllum oldhamii* (G-I) and *Schima superba* (J-L) and the metabolite diversity in the exudates, roots, and leaves (calculated as Shannon diversity). The diversity levels were the diversity level of the local neighbourhood of a single tree. Dotted lines indicate the confidence interval of the regression (standard deviation x 3). N is the number of replicates, r is the correlation coefficient and p the p value

**Table S1:** Theoretical plot diversity vs. average local diversity of the target trees in the plots. The number of species in the direct local neighbourhood (10 adjacent grid squares) of the tree species pair the target trees belonged to. The differential between plot and local diversity can be partially explained by the fact that the latter is maximized to 11 (1 pair neighbour plus 10 grid squares around). Moreover, some grid cells were empty due to tree mortality or occupied by a different species than expected due to misplanting or invasion of empty grid cells by the nearest neighbouring species.

| Plot Diversity | Average Local Diversity |
|---|---|
| 1 | 1.08 |
| 2 | 2.22 |
| 4 | 3.80 |
| 8 | 5.11 |
| 16 | 4.00 |
| 24 | 5.33 |

**Table S2:** Mass features that have the highest positive loading on the principal component PC1 as well as the highest positive or negative loading on PC2 of the principal component analysis of the metabolome analysis of the exudates of the four tree species *C. camphora, C. glauca, D. oldhamii, and S. superba*. Putative annotations were achieved by either: a manual interpretation using sum formula, literature data, spectral similarity and fragmentation pattern b computing the sum formula, c comparison to spectral libraries, d comparison to an in-house spectral library of manually assigned species specific features.

| Loadings | | RT [min] | m/z measured | Molecular Mass | Adduct type | Molecular Formula | Putative Annotation | |
|---|---|---|---|---|---|---|---|---|
| PC1 | PC2 | | | | | | | |
| 0.124 | 0.013 | 10.96 | 514.39155 | 1026.7685 | $[M+H+H]^{2+}$ | $C_{68}H_{102}N_2O_5$ | D. oldhamii alkaloid | b,d |
| 0.12 | 0.014 | 10.98 | 515.39248 | 514.3852 | $[M+H]^+$ | | D. oldhamii alkaloid, isotope peak | a |
| 0.121 | 0.012 | 11.46 | 470.36211 | 469.35483 | $[M+H]^+$ | $C_{30}H_{47}NO_3$ | Secodaphniphylline/ Daphnioldhanin D | b,d, |
| 0.125 | 0.015 | 11.64 | 468.34629 | 467.33901 | $[M+H]^+$ | $C_{30}H_{45}NO_3$ | D. oldhamii alkaloid similar to Secodaphniphylline/Daphnioldhanin | b,d, |
| 0.14 | 0.014 | 11.87 | 470.36367 | 938.7128 | $[M+H+H]^{2+}$ | $C_{68}H_{92}NO$ | similar to Secodaphniphylline/ Daphnioldhanin D | b,d, |
| 0.13 | 0.016 | 11.89 | 471.36615 | 470.35888 | $[M+H]^+$ | | similar to Secodaphniphylline/ Daphnioldhanin D, isotope peak | d, |
| 0.118 | 0.013 | 11.60 | 472.37723 | 471.36996 | $[M+H]^+$ | $C_{30}H_{49}NO_3$ | unknown | b |
| -0.038 | 0.109 | 4.17 | 314.13818 | 313.13091 | $[M+H]^+$ | $C_{18}H_{19}NO_4$ | Laurolitsine-like | b,d, |
| -0.028 | 0.112 | 12.11 | 133.10099 | 132.09371 | $[M+H]^+$ | $C_{10}H_{12}$ | C. camphora terpenoid, fragment | b,d |
| -0.03 | 0.1095 | 12.11 | 219.17428 | 218.167 | $[M+H]^+$ | $C_{15}H_{22}O$ | C. camphora terpenoid | b,d |
| -0.028 | 0.118 | 12.12 | 175.14823 | 174.14096 | $[M+H]^+$ | $C_{13}H_{18}$ | C. camphora terpenoid, fragment | b,d |
| -0.032 | 0.145 | 12.53 | 189.16376 | 188.15648 | $[M+H]^+$ | $C_{14}H_{20}$ | C. camphora terpenoid, fragment | b,d |
| -0.03 | 0.15 | 12.76 | 251.1633 | 250.15664 | [M+H]+, | $C_{15}H_{22}O_3$ | C. camphora benzochinone | c |
| -0.024 | 0.1 | 13.06 | 233.15344 | 250.15627 | $[M+H-H_2O]^+$, | $C_{15}H_{22}O_3$ | C. camphora phenylpropanoid | c |

| | | | | | | | | |
|---|---|---|---|---|---|---|---|---|
| -0.02 | -0.046 | 3.79 | 435.05883 | 434.05134 | $[M+H]^+$ | $C_{19}H_{14}O_{12}$ | S. superba, coumarin? | c |
| -0.02 | -0.054 | 10.74 | 1137.56907 | 1136.5618 | $[M+H]^+$ | $C_{54}H_{88}O_{25}$ | S. superba saponine | b,d |
| -0.018 | -0.046 | 11.37 | 1121.57357 | 1120.5658 | $[M+H]^+$ | $C_{54}H_{88}O_{24}$ | S. superba saponine | b,d |
| -0.02 | -0.052 | 11.6 | 1119.55785 | 1118.55 | $[M+H]^+$ | $C_{54}H_{86}O_{24}$ | S. superba saponine | b,d |
| -0.02 | -0.052 | 11.7 | 1105.5782 | 1104.5697 | $[M+H]^+$ | $C_{54}H_{88}O_{23}$ | S. superba saponine | b,d |

**Table S3:** Mass features that have the highest positive loading on the principal component PC1 as well as the highest positive or negative loading on PC2 of the principal component analysis of the metabolome analysis of the roots of the four tree species *C. camphora, C. glauca, D. oldhamii, and S. superba*. Putative annotations were achieved by either: a manual interpretation using sum formula, literature data, spectral similarity and fragmentation pattern, b computing the sum formula, c comparison to spectral libraries, d comparison to an in-house spectral library of manually assigned species specific features.

| Loadings | | RT | m/z | Molecular | Adduct | Molecular | Putative Annotation | |
|---|---|---|---|---|---|---|---|---|
| PC1 | PC2 | [min] | measured | Mass | type | Formula | | |
| 0.15 | 0.009 | 10.39 | 372.25269 | 371.24541 | $[M+H]^+$ | $C_{23}H_{33}NO_3$ | D. oldhamii alkaloid similar to Oldhamine A | b,d |
| 0.149 | 0.013 | 10.98 | 515.39248 | 514.3852 | $[M+H]^+$ | $C_{35}H_{49}N_2O$ | D. oldhamii alkaloid, isotope peak | a |
| 0.145 | 0.007 | 11.46 | 470.36211 | 469.35483 | $[M+H]^+$ | $C_{30}H_{47}NO_3$ | Secodaphniphylline/ Daphnioldhanin D | b,d |
| 0.143 | 0.014 | 11.64 | 492.27394 | 491.26666 | $[M+H]^+$ | $C_{30}H_{37}NO_5$ | similar to cevadine | c |
| 0.144 | 0.006 | 11.87 | 470.36367 | 938.7128 | $[M+H+H]_2+$ | $C_{68}H_{92}NO$ | D. oldhamii alkaloid | b,d |
| 0.152 | 0.013 | 11.89 | 471.36615 | 470.35888 | $[M+H]^+$ | | D. oldhamii alkaloid, isotope peak | a |
| -0.039 | 0.165 | 3.9 | 355.10194 | 354.09455 | $[M+H]^+$ | $C_{16}H_{18}O_9$ | S. superba,phenylpropanoid? | c |
| -0.024 | 0.105 | 11.6 | 1119.55785 | 1118.55 | $[M+H]^+$ | $C_{54}H_{86}O_{24}$ | S. superba saponine | b,d |
| -0.028 | 0.12 | 11.7 | 1105.5782 | 1104.5697 | $[M+H]^+$ | $C_{54}H_{88}O_{23}$ | S. superba saponine | b,d |
| -0.022 | 0.104 | 12.29 | 1219.61076 | 1218.6035 | $[M+H]^+$ | $C_{59}H_{94}O_{26}$ | S. superba saponine | b,d |
| -0.035 | 0.164 | 12.71 | 1103.56302 | 1102.5557 | $[M+H]^+$ | $C_{54}H_{86}O_{23}$ | S. superba saponine | b,d |
| -0.03 | -0.17 | 4.17 | 314.13818 | 313.13091 | $[M+H]^+$ | $C_{18}H_{19}NO_4$ | Laurolitsine-like | b,d |
| -0.025 | -0.14 | 4.18 | 297.1116 | 296.10432 | $[M+H]^+$ | $C_{18}H_{16}O_4$ | Laurolitsine-like, fragment -NH3 | a,d |
| -0.02 | -0.105 | 4.32 | 865.19718 | 864.1899 | $[M+H]^+$ | $C_{45}H_{36}O_{18}$ | C. camphora Procyanidin | c |
| -0.028 | -0.165 | 4.5 | 330.1693 | 329.16203 | $[M+H]^+$ | $C_{19}H_{23}NO_4$ | (S)-Reticuline | a |
| -0.015 | -0.097 | 13.44 | 138.06744 | 137.06016 | $[M+H]^+$ | $C_8H_9O_2$ | unknown | b |
| -0.01 | -0.086 | 15.85 | 205.195 | 204.18772 | $[M+H]^+$ | $C_{15}H_{24}$ | C. camphora sesquiterpene | b,d |

**Table S4:** Mass features that have the highest positive loading on the principal component PC1 as well as the highest positive or negative loading on PC2 of the principal component analysis of the metabolome analysis of the leaves of the four tree species *C. camphora, C. glauca, D. oldhamii, and S. superba*. Putative annotations were achieved by either: **(a)** manual interpretation using sum formula, literature data, spectral similarity and fragmentation pattern, **(b)** computing the sum formula, **(c)** comparison to spectral libraries, **(d)** comparison to an in-house spectral library of manually assigned species specific features.

| **Loadings** | | **RT [min]** | **m/z measured** | **Molecular Mass** | **Adduct type** | **Molecular Formula** | **Putative Annotation** | |
|---|---|---|---|---|---|---|---|---|
| **PC1** | **PC2** | | | | | | | |
| 0.139 | 0.01 | 0.99 | 163.06022 | 162.05294 | $[M+H]^+$ | $C_6H_{10}O_5$ | Disaccharide fragment | a,c |
| 0.135 | -0.005 | 0.99 | 325.1125 | 324.10467 | $[M+H]^+$ | $C_{12}H_{20}O_{10}$ | Disaccharide | a,c |
| 0.18 | -0.015 | 4.2 | 175.03915 | 192.04235 | $[M+H-H_2O]^+$ | $C_{10}H_8O_4$ | 7,8-Dihydroxy-4-methylcoumarin, fragment of C12H10O5 | a,c |
| 0.14 | -0.01 | 6.72 | 579.16969 | 578.16241 | $[M+H]^+$ | $C_{27}H_{30}O_{14}$ | D. oldhamii flavonoid glycoside, | a,c |
| 0.131 | 0.01 | 11.14 | 271.05972 | 270.05244 | $[M+H]^+$ | $C_{21}H_{20}O_{10}$ | D. oldhamii flavonoid? | c |
| 0.129 | -0.01 | 17.45 | 425.37556 | 424.36854 | $[M+H]^+$ | $C_{30}H_{48}O$ | Lupenone | c |
| -0.02 | 0.105 | 13.31 | 353.13754 | 352.13035 | $[M+H]^+$ | $C_{21}H_{20}O_5$ | unknown | b |
| -0.04 | 0.15 | 15.62 | 203.17948 | 202.1722 | $[M+H]^+$ | $C_{15}H_{22}$ | C. camphora sesquiterpene | b,d |
| -0.025 | 0.09 | 16.18 | 205.195 | 204.18773 | $[M+H]^+$ | $C_{15}H_{24}$ | C. camphora sesquiterpene | b,d |
| -0.02 | 0.12 | 17.46 | 609.27081 | 608.26353 | $[M+H]^+$ | $C_{35}H_{36}N_4O_6$ | similar to Reserpine | c |
| -0.006 | 0.095 | 17.7 | 594.27959 | 593.27232 | $[M+H]^+$ | $C_{19}H_{43}N_6Na_2O_{12}$ | unknown | b |
| -0.025 | -0.07 | 1.02 | 193.07078 | 192.06346 | $[M+H]^+$ | $C_7H_{12}O_6$ | Quinic acid | c |
| -0.038 | -0.05 | 10.38 | 741.20234 | 740.19507 | $[M+H]^+$ | $C_{36}H_{36}O_{17}$ | C. glauca similar to Proanthocyanidin A | b,c |
| -0.05 | -0.08 | 12.64 | 1037.65606 | 1036.6486 | $[M+H]^+$ | $C_{60}H_{92}O_{14}$ | C. glauca terpenoid dimer | b,d |
| -0.04 | -0.065 | 12.66 | 501.32039 | 500.31311 | $[M+H]^+$ | $C_{30}H_{44}O_6$ | C. glauca terpenoid fragment | b,d |
| -0.04 | -0.06 | 12.89 | 825.20265 | 824.19537 | $[M+H]^+$ | $C_{43}H_{36}O_{17}$ | C. glauca polyphenol 12.89min : 824.19537m/ | b,c |
| -0.035 | -0.052 | 13.08 | 825.20264 | 824.19536 | $[M+H]^+$ | $C_{43}H_{36}O_{17}$ | C. glauca polyphenol 13.08min : 824.19536m/z | b,c |

**Table S5:** Results of the comparison of the metabolome composition in exudates, roots, and leaves of the four different tree species *Cinnamomum camphora, Cyclobalanopsis glauca, Daphniphyllum oldhamii, and Schima superba* across the different levels of diversity of the experimental plot by multiple response permutation procedures. P - values under the different sections are the global values, while values in the sections are the p-values of the pairwise comparisons. Significant p-values smaller than 0.05 are in bold.

| | *Cinnamomum camphora* | | | | *Cyclobalanopsis glauca* | | | | *Daphniphyllum oldhamii* | | | | *Schima superba* | | | |
|---|---|---|---|---|---|---|---|---|---|---|---|---|---|---|---|---|
| **Exudates** | | | | | | | | | | | | | | | | |
| | 2 | 4 | 8 | 16 | 2 | 4 | 8 /16 | | 2 | 4 | 8 | 16 | 2 | 4 | 8 | 16 |
| 1 | - | - | - | - | 0.57 | 0.49 | 0.53 | | 0.42 | 0.54 | 0.57 | 0.52 | 0.47 | 0.52 | 0.55 | 0.50 |
| 2 | | - | - | - | | 0.16 | 0.36 | | | 0.44 | 0.47 | 0.50 | | 0.45 | 0.47 | 0.45 |
| 4 | | | - | - | | | 0.50 | | | | 0.51 | 0.36 | | | 0.53 | 0.47 |
| 8 | | | | - | | | | | | | | 0.52 | | | | 0.44 |
| | p = 0.535 | | | | **p = 0.018** | | | | **p = 0.006** | | | | **p = 0.001** | | | |
| **Roots** | | | | | | | | | | | | | | | | |
| | 2 | 4 | 8 | 16 | 2 | 4 | 8 | 16 | 2 | 4 | 8 | 16 | 2 | 4 | 8 | 16 |
| 1 | - | - | - | - | - | - | - | - | 0.47 | 0.48 | 0.47 | 0.57 | - | - | - | - |
| 2 | | - | - | - | | - | - | - | | 0.57 | 0.41 | 0.51 | | - | - | - |
| 4 | | | - | - | | | - | - | | | 0.54 | 0.52 | | | - | - |
| 8 | | | | - | | | | - | | | | 0.52 | | | | - |
| | p = 0.279 | | | | p = 0.838 | | | | **p = 0.008** | | | | p = 0.204 | | | |
| **Leaves** | | | | | | | | | | | | | | | | |
| | 2 | 4 | 8 | 16 | 2 | 4 | 8 | 16 | 2 | 4 | 8 | 16 | 2 | 4 | 8 | 16 |
| 1 | 0.54 | 0.43 | 0.52 | 0.49 | 0.57 | 0.49 | 0.53 | 0.62 | - | - | - | - | 0.49 | 0.48 | 0.44 | 0.45 |
| 2 | | 0.50 | 0.48 | 0.43 | | 0.16 | 0.36 | 0.49 | | - | - | - | | 0.51 | 0.60 | 0.55 |
| 4 | | | 0.53 | 0.52 | | | 0.50 | 0.59 | | | - | - | | | 0.34 | 0.38 |
| 8 | | | | 0.46 | | | | 0.55 | | | | - | | | | 0.57 |
| | **p = 0.002** | | | | **p = 0.013** | | | | p = 0.081 | | | | **p = 0.027** | | | |

**Table S6:** Results of the comparison of the metabolome composition in exudates, roots, and leaves of the four different tree species *Cinnamomum camphora, Cyclobalanopsis glauca, Daphniphyllum oldhamii, and Schima superba* across the different levels of local diversity by multiple response permutation procedures. P - values under the different sections are the global values, while values in the sections are the p-values of the pairwise comparisons. Significant p-values smaller than 0.05 are in bold.

| | *Cinnamomum camphora* | | | *Cyclobalanopsis glauca* | | | *Daphniphyllum oldhamii* | | *Schima superba* | | |
|---|---|---|---|---|---|---|---|---|---|---|---|
| **Exudates** | | | | | | | | | | | |
| | 2 | 3 /4 | 5/6/7 | 2 | 3/4/5 | | 3 /4 | 5 /6 | 2 | 3/4 | 5 /6 |
| 1 | - | - | - | - | - | | - | - | 0.360 | 0.533 | 0.515 |
| 2 | | - | - | | - | | | - | | 0.310 | 0.290 |
| 3/4 | | | - | | | | | | | | 0.385 |
| | p = 0.817 | | | p = 0.092 | | | p =0.846 | | **p = 0.040** | | |
| **Roots** | | | | | | | | | | | |
| | 2 | 3/ 4 | 5/6/7 | 2 | 3/4 | 5/6/7 | 3 /4 | 5 /6 | 2 | 3 /4 | 5/6 |
| 1 | - | - | - | - | - | - | - | - | - | - | - |
| 2 | | - | - | | - | - | | - | | - | - |
| 3/4 | | | - | | | - | | | | | - |
| | p = 0.842 | | | p = 0.873 | | | p =0.846 | | p = 0.169 | | |
| **Leaves** | | | | | | | | | | | |
| | 2 | 3 /4 | 5 /6/7 | 2 | 3/4/5 | | 3 /4 | 5 /6 | 2 | 3 /4 | 5/6 |
| 1 | 0.46 | 0.45 | 0.44 | 0.516 | 0.449 | | - | - | 0.50 | 0.51 | 0.56 |
| 2 | | 0.47 | 0.51 | | 0.539 | | | - | | 0.55 | 0.57 |
| 3/4 | | | 0.53 | | | | | | | | 0.57 |
| | **p = 0.001** | | | **p = 0.013** | | | p =0.846 | | **p = 0.016** | | |

**Table S7:** Percentage of the total intensities of the m/z features that could be classified by the ClassyFire tool

| | *Cinnamomum camphora* | *Cyclobalanopsis glauca* | *Daphniphyllum oldhamii* | *Schima superba* |
|---|---|---|---|---|
| Exudates | 37.1% | 29.2% | 40.6% | 30.2% |
| Roots | 40.9% | 22.5% | 38.0% | 31.7% |
| Leaves | 35.4% | 27.8% | 46.0% | 32.4% |

**Table S8:** Top 20 VIPs (Variable importance in projection) of the PLS-DA analysis of the exudates. RT is the retention time, m/z the measured mass to charge ratio and Comp1 the VIP coefficient on the first component of the PLS-DA

| ***Cinnamomum camphora*** | | | | | | ***Cyclobalanopsis glauca*** | | | | | |
|---|---|---|---|---|---|---|---|---|---|---|---|
| Local Diversity | | | Plot Diversity | | | Local Diversity | | | Plot Diversity | | |
| RT in min | m/z | Comp. 1 | RT in min | m/z | Comp. 1 | RT in min | m/z | Comp. 1 | RT in min | m/z | Comp. 1 |
| 1.42 | 228.10436 | 4.8187 | 1.42 | 202.18024 | 5.0764 | 1.42 | 202.18024 | 7.1044 | 1.04 | 188.03504 | 6.0632 |
| 3.43 | 153.12735 | 5.2372 | 4.02 | 314.13819 | 5.5816 | 3.53 | 227.17516 | 5.4218 | 1.09 | 148.04242 | 6.8564 |
| 3.94 | 227.17537 | 4.668 | 4.3 | 328.15369 | 5.6579 | 3.94 | 227.17537 | 14.526 | 3.94 | 227.17537 | 17.368 |
| 4.3 | 328.15369 | 9.5896 | 4.6 | 179.0703 | 5.0369 | 4.91 | 170.1182 | 5.1506 | 5.12 | 453.34251 | 5.5853 |
| 4.37 | 329.15842 | 5.0414 | 4.87 | 181.04965 | 8.3677 | 5.12 | 453.3425 | 7.7029 | 5.59 | 453.34282 | 11.978 |
| 4.42 | 342.16923 | 5.2369 | 11.1 | 237.18458 | 5.8093 | 5.6 | 453.34282 | 7.0262 | 10.75 | 212.16459 | 8.1887 |
| 4.5 | 330.1693 | 4.8708 | 11.14 | 219.17426 | 5.406 | 5.95 | 566.42667 | 6.692 | 10.99 | 514.38869 | 6.5341 |
| 4.87 | 181.04965 | 4.9672 | 11.22 | 267.15853 | 5.594 | 10.75 | 212.16459 | 7.6176 | 11.12 | 246.24259 | 8.8017 |
| 5.35 | 284.12747 | 4.7312 | 11.5 | 226.18004 | 6.4013 | 11.12 | 246.2426 | 8.9829 | 11.5 | 226.18004 | 14.631 |
| 6.15 | 179.06499 | 4.8949 | 11.95 | 249.14829 | 5.3605 | 11.5 | 226.18004 | 15.763 | 11.5 | 244.19053 | 6.6429 |
| 11.95 | 249.14829 | 5.055 | 12.01 | 189.16381 | 6.522 | 11.5 | 244.19054 | 7.0947 | 11.64 | 492.27394 | 6.7484 |
| 12.01 | 189.16381 | 6.8297 | 12.11 | 219.17428 | 7.2144 | 11.64 | 492.27394 | 6.4472 | 11.85 | 512.37304 | 7.2396 |
| 12.11 | 219.17428 | 6.6506 | 12.17 | 274.27352 | 13.43 | 11.85 | 512.37304 | 5.6703 | 11.91 | 470.36206 | 6.0378 |
| 12.17 | 274.27352 | 9.6955 | 12.29 | 203.17939 | 5.8134 | 11.91 | 470.36206 | 5.9034 | 12.21 | 192.13832 | 9.5513 |
| 12.29 | 203.17939 | 5.2158 | 12.49 | 235.16907 | 6.1672 | 12.21 | 192.1386 | 7.1434 | 12.21 | 192.1386 | 6.8414 |
| 13.38 | 189.1638 | 4.5694 | 12.7 | 205.15859 | 4.8727 | 12.21 | 192.13832 | 7.1352 | 12.22 | 318.2999 | 10.542 |
| 13.39 | 235.169 | 6.5208 | 13.38 | 189.1638 | 6.6552 | 12.22 | 318.2999 | 8.8347 | 12.3 | 362.3258 | 6.0485 |
| 13.55 | 121.10109 | 6.0195 | 13.55 | 121.10109 | 8.5737 | 12.47 | 288.28911 | 8.0547 | 12.3 | 288.28942 | 5.9189 |
| 13.56 | 219.17429 | 5.2932 | 13.56 | 219.17428 | 6.1472 | 14.72 | 291.19456 | 6.3917 | 12.47 | 288.2891 | 10.739 |
| 13.93 | 180.13865 | 5.3552 | 13.75 | 415.21091 | 5.9697 | 14.76 | 200.20087 | 7.7015 | 13.87 | 251.1629 | 6.4872 |

**Table S8:** continued

| *Daphniphyllum oldhamii* | | | | | | *Schima superba* | | | | | |
|---|---|---|---|---|---|---|---|---|---|---|---|
| Local Diversity | | | Plot Diversity | | | Local Diversity | | | Plot Diversity | | |
| RT in min | mz | Comp. 1 | RT in min | m/z | Comp. 1 | RT in min | m/z | Comp. 1 | RT in min | m/z | Comp. 1 |
| 4.05 | 370.16405 | 6.0586 | 3.94 | 227.17537 | 9.0881 | 1.09 | 148.04242 | 6.0396 | 1.42 | 202.18024 | 7.0498 |
| 4.52 | 358.23714 | 10.367 | 4.52 | 358.23714 | 7.4647 | 1.42 | 202.18024 | 7.1942 | 3.9 | 355.10193 | 5.2054 |
| 4.54 | 354.16941 | 5.9062 | 4.54 | 354.16941 | 5.9403 | 1.42 | 228.10436 | 6.9697 | 3.94 | 227.17537 | 11.04 |
| 6.64 | 362.26819 | 5.3018 | 4.65 | 358.23712 | 6.4625 | 3.94 | 227.17537 | 9.0478 | 4.48 | 223.02377 | 5.5134 |
| 8.69 | 430.25933 | 5.4872 | 5.01 | 342.24196 | 9.1282 | 4.54 | 354.16941 | 5.6372 | 10.74 | 1137.5691 | 6.5988 |
| 8.99 | 346.27328 | 6.5654 | 5.68 | 368.18489 | 9.2146 | 5.6 | 453.34282 | 6.1941 | 11.03 | 1121.5737 | 5.0269 |
| 10.39 | 372.25269 | 11.735 | 7.29 | 346.2733 | 6.9249 | 10.74 | 1137.5691 | 6.9254 | 12.17 | 274.27352 | 16.867 |
| 10.73 | 486.35682 | 6.7895 | 10.17 | 490.38813 | 6.4826 | 11.64 | 492.27394 | 5.7214 | 12.21 | 192.1386 | 13.104 |
| 10.87 | 530.38303 | 7.1327 | 10.4 | 372.25269 | 9.0142 | 12.17 | 274.27352 | 9.4563 | 12.23 | 290.26861 | 5.2269 |
| 11.27 | 360.28908 | 9.1453 | 10.6 | 376.28374 | 6.3494 | 12.21 | 192.1386 | 11.091 | 14.72 | 291.19456 | 5.7357 |
| 11.38 | 360.28911 | 6.2674 | 11.31 | 544.36273 | 8.1501 | 14.14 | 473.36151 | 7.5596 | 14.87 | 291.25241 | 5.2537 |
| 11.64 | 492.27394 | 7.3228 | 11.61 | 493.27773 | 11.053 | 15.08 | 309.20543 | 5.639 | 15.08 | 309.20543 | 5.8408 |
| 11.66 | 544.36264 | 7.012 | 11.66 | 544.36264 | 11.343 | 16.32 | 228.23199 | 6.6841 | 16.62 | 688.52008 | 6.7713 |
| 11.73 | 526.35183 | 5.6858 | 11.73 | 526.35183 | 6.0801 | 16.62 | 688.52008 | 5.7835 | 16.65 | 644.49392 | 7.3306 |
| 11.8 | 529.37158 | 11.385 | 11.8 | 529.37158 | 9.3136 | 16.65 | 644.49393 | 6.4211 | 16.65 | 627.46665 | 5.672 |
| 11.82 | 528.36772 | 5.2706 | 11.82 | 528.36772 | 6.9499 | 16.69 | 621.39715 | 6.0536 | 16.69 | 621.39715 | 5.9187 |
| 11.92 | 476.27741 | 7.2155 | 11.87 | 357.10221 | 6.8664 | 16.72 | 556.44146 | 7.3728 | 16.72 | 556.44146 | 7.7387 |
| 11.97 | 526.35162 | 8.6556 | 11.96 | 478.2938 | 6.0834 | 16.75 | 512.41509 | 6.9598 | 16.75 | 512.41509 | 6.884 |
| 12.07 | 529.37084 | 7.0534 | 12.22 | 318.2999 | 6.2992 | 16.78 | 451.36219 | 6.1006 | 16.78 | 451.36219 | 6.2785 |
| 12.23 | 290.26861 | 6.2781 | 12.58 | 514.38834 | 7.6784 | 16.81 | 407.33586 | 5.9881 | 16.81 | 407.33586 | 6.2031 |

**Table S9:** Top 20 VIPs (Variable importance in projection) of the PLS-DA analysis of the roots. RT is the retention time, m/z the measured mass to charge ratio and Comp1 the VIP coefficient on the first component of the PLS-DA

| ***Cinnamomum camphora*** | | | | | | ***Cyclobalanopsis glauca*** | | | | | |
|---|---|---|---|---|---|---|---|---|---|---|---|
| Local Diversity | | | Plot Diversity | | | Local Diversity | | | Plot Diversity | | |
| RT in min | m/z | Comp. 1 | RT in min | m/z | Comp. 1 | RT in min | m/z | Comp. 1 | RT in min | m/z | Comp. 1 |
| 4.3 | 328.1537 | 10.157 | 13.56 | 219.1743 | 11.623 | 17.18 | 439.3572 | 14.191 | 17.18 | 439.3572 | 12.4 |
| 13.44 | 138.0674 | 9.6991 | 16.15 | 219.1742 | 10.097 | 10.64 | 473.3256 | 6.495 | 14.76 | 200.2009 | 8.113 |
| 16.15 | 219.1742 | 8.1729 | 15.85 | 205.195 | 9.6782 | 1.04 | 952.105 | 6.1425 | 10.64 | 473.3256 | 7.3091 |
| 17.43 | 439.3569 | 8.0761 | 15.99 | 123.1168 | 7.8451 | 1.03 | 1084.148 | 5.4862 | 10.63 | 455.315 | 6.1942 |
| 13.41 | 179.1066 | 7.9284 | 5.35 | 284.1275 | 7.6158 | 10.64 | 455.315 | 5.4739 | 10.64 | 501.3208 | 6.1242 |
| 4.59 | 359.132 | 7.8158 | 14.76 | 200.2009 | 7.3405 | 10.64 | 501.3208 | 5.446 | 10.7 | 1037.655 | 6.0326 |
| 4.61 | 135.044 | 7.7722 | 12.76 | 251.1633 | 6.9258 | 17.43 | 439.3569 | 5.3926 | 10.3 | 665.3888 | 5.1712 |
| 4.6 | 179.0703 | 7.7399 | 15.43 | 219.1743 | 6.8844 | 10.3 | 665.3888 | 5.3011 | 1.04 | 952.105 | 5.0217 |
| 13.39 | 164.0831 | 6.8816 | 4.01 | 291.0858 | 6.7514 | 14.76 | 200.2009 | 5.1525 | 12.58 | 501.3207 | 4.5484 |
| 15.99 | 123.1168 | 6.2 | 4.17 | 314.1382 | 6.0043 | 10.7 | 1037.656 | 4.7216 | 17.24 | 439.3575 | 4.5335 |
| 4.01 | 291.0858 | 5.9979 | 4.42 | 291.0858 | 5.904 | 12.76 | 671.342 | 4.6068 | 12.76 | 671.342 | 4.4831 |
| 15.85 | 205.195 | 5.9746 | 4.32 | 865.1972 | 5.7035 | 17.95 | 352.3387 | 4.3314 | 16.61 | 453.336 | 4.4712 |
| 15.43 | 219.1743 | 5.5438 | 3.66 | 579.1494 | 5.521 | 16.29 | 455.3511 | 4.2736 | 16.34 | 409.3459 | 4.2858 |
| 4.59 | 717.2596 | 5.4091 | 4.01 | 139.0389 | 5.2603 | 1 | 187.0588 | 4.2615 | 10.63 | 519.3309 | 4.2796 |
| 17.32 | 352.3386 | 5.2873 | 17.24 | 311.2563 | 5.2408 | 10.65 | 698.4104 | 4.2419 | 1.03 | 1084.148 | 4.1901 |
| 4.42 | 291.0858 | 5.2684 | 15.42 | 235.169 | 5.1612 | 10.32 | 473.3252 | 4.2113 | 16.29 | 455.3511 | 4.1574 |
| 0.99 | 145.0495 | 5.1209 | 13.44 | 138.0674 | 5.0317 | 12.58 | 501.3206 | 4.0807 | 10.32 | 473.3252 | 4.1256 |
| 4.32 | 865.1972 | 4.9384 | 4.18 | 297.1116 | 4.9137 | 3.27 | 935.079 | 4.0596 | 17.92 | 455.3509 | 3.9269 |
| 1.04 | 130.0863 | 4.9334 | 4.15 | 579.1495 | 4.8836 | 4.76 | 435.0557 | 3.9164 | 17.43 | 439.3569 | 3.916 |
| 1 | 846.3081 | 4.8824 | 16 | 135.1167 | 4.8674 | 10.63 | 519.3309 | 3.8954 | 10.65 | 698.4104 | 3.9136 |

**Table S9:** continued

| *Daphniphyllum oldhamii* | | | | | | *Schima superba* | | | | | |
|---|---|---|---|---|---|---|---|---|---|---|---|
| Local Diversity | | | Plot Diversity | | | Local Diversity | | | Plot Diversity | | |
| RT in min | m/z | Comp. 1 | RT in min | m/z | Comp. 1 | RT in min | m/z | Comp. 1 | RT in min | m/z | Comp. 1 |
| 17.24 | 439.3575 | 12.992 | 4.13 | 360.2526 | 10.912 | 17.24 | 439.3575 | 11.975 | 17.18 | 439.3572 | 17.516 |
| 11.92 | 476.2774 | 11.568 | 4.52 | 358.2371 | 8.9585 | 4.11 | 355.1019 | 9.5369 | 17.24 | 439.3575 | 11.621 |
| 4.13 | 360.2526 | 10.905 | 4.65 | 358.2371 | 9.4304 | 17.43 | 439.3569 | 8.6519 | 17.43 | 439.3569 | 8.0153 |
| 4.65 | 358.2371 | 9.8866 | 7.98 | 346.2727 | 7.1706 | 14.76 | 200.2009 | 8.2216 | 10.74 | 1137.569 | 7.5572 |
| 0.99 | 163.0602 | 8.3065 | 8.59 | 430.2591 | 7.0268 | 16.6 | 471.3466 | 8.0546 | 16.6 | 471.3466 | 7.2888 |
| 11.15 | 532.3989 | 8.1384 | 10.17 | 490.3881 | 7.9874 | 3.9 | 355.1019 | 7.2651 | 4.11 | 355.1019 | 6.7619 |
| 0.99 | 145.0495 | 8.0415 | 11.08 | 360.2888 | 7.7217 | 3.19 | 371.0956 | 7.1043 | 425.3403 | 425.3403 | 5.1262 |
| 17.43 | 439.3569 | 7.9489 | 11.38 | 360.2891 | 8.3107 | 10.74 | 1137.569 | 6.4723 | 3.41 | 435.0588 | 5.1156 |
| 11.64 | 468.3463 | 7.8519 | 11.42 | 472.3765 | 7.1401 | 15.02 | 453.3352 | 5.7705 | 17.32 | 352.3386 | 5.0503 |
| 12.58 | 514.3883 | 7.4573 | 11.6 | 472.3772 | 7.8424 | 17.76 | 425.3403 | 5.6483 | 17.95 | 352.3387 | 5.0495 |
| 3.11 | 195.0653 | 6.508 | 11.64 | 468.3463 | 7.3776 | 16.59 | 219.1742 | 5.4541 | 16.73 | 553.3892 | 4.9317 |
| 5.01 | 342.242 | 6.1976 | 11.8 | 529.3716 | 6.3484 | 13.34 | 437.339 | 5.2523 | 0.99 | 145.0495 | 4.915 |
| 11.38 | 360.2891 | 6.077 | 11.82 | 528.3677 | 9.593 | 13.77 | 437.3403 | 5.1655 | 16.6 | 219.1742 | 4.8407 |
| 8.04 | 344.2575 | 5.9888 | 11.84 | 513.3769 | 8.9911 | 17.32 | 352.3386 | 5.0886 | 0.97 | 324.1646 | 4.7096 |
| 3.11 | 749.2498 | 5.9473 | 11.92 | 476.2774 | 11.537 | 16.61 | 453.336 | 5.0342 | 14.76 | 200.2009 | 4.5456 |
| 4.52 | 358.2371 | 5.8529 | 11.96 | 478.2938 | 9.2695 | 16.53 | 472.3496 | 5.0281 | 13.34 | 437.339 | 4.4341 |
| 4.95 | 388.2475 | 5.8288 | 12.38 | 504.3087 | 8.8074 | 17.95 | 352.3387 | 4.8355 | 13.77 | 437.3403 | 4.3557 |
| 12.33 | 502.2938 | 5.7148 | 12.58 | 514.3883 | 6.1308 | 0.99 | 163.0602 | 4.6749 | 11.37 | 1121.574 | 4.1129 |
| 11.6 | 472.3772 | 5.4952 | 17.24 | 439.3575 | 19.842 | 5.52 | 331.1534 | 4.6668 | 15.02 | 453.3352 | 4.0696 |
| 16.34 | 409.3459 | 5.4522 | 17.43 | 439.3569 | 10.547 | 0.96 | 407.1654 | 4.6568 | 12.75 | 1101.547 | 3.9945 |

**Table S10:** Top 20 VIPs (Variable importance in projection) of the PLS-DA analysis of the leaves. RT is the retention time, m/z the measured mass to charge ratio and Comp1 the VIP coefficient on the first component of the PLS-DA

| ***Cinnamomum camphora*** | | | | | | ***Cyclobalanopsis glauca*** | | | | | |
|---|---|---|---|---|---|---|---|---|---|---|---|
| Local Diversity | | | Plot Diversity | | | Local Diversity | | | Plot Diversity | | |
| RT in min | m/z | Comp. 1 | RT in min | m/z | Comp. 1 | RT in min | m/z | Comp. 1 | RT in min | m/z | Comp. 1 |
| 15.62 | 203.1795 | 22.286 | 13.55 | 121.1013 | 16.528 | 17.39 | 414.3571 | 14.014 | 17.39 | 414.3571 | 21.24 |
| 13.56 | 219.1743 | 13.834 | 17.7 | 593.276 | 13.779 | 5.19 | 611.1606 | 8.3792 | 17.46 | 609.2708 | 9.537 |
| 17.7 | 594.2796 | 12.047 | 17.7 | 594.2796 | 13.422 | 12.64 | 1037.656 | 8.2708 | 17.65 | 621.2707 | 8.2628 |
| 17.46 | 609.2708 | 10.947 | 13.56 | 219.1743 | 13.013 | 17.33 | 171.1382 | 6.7899 | 17.25 | 609.2706 | 8.1209 |
| 15.62 | 147.1168 | 10.229 | 17.46 | 609.2708 | 11.755 | 14.51 | 277.2157 | 6.1879 | 17.18 | 439.3572 | 7.1891 |
| 17.7 | 593.276 | 9.8349 | 16.18 | 205.195 | 11.538 | 17.65 | 621.2707 | 6.1709 | 17.33 | 171.1382 | 7.113 |
| 6.03 | 595.1657 | 8.0823 | 15.62 | 203.1795 | 10.871 | 17.43 | 439.3569 | 5.4976 | 5.19 | 611.1606 | 6.5255 |
| 6.27 | 625.176 | 7.8973 | 17.25 | 609.2706 | 8.5757 | 12.58 | 501.3206 | 5.0305 | 12.58 | 501.3207 | 6.3277 |
| 15.47 | 203.1795 | 7.3736 | 16.17 | 121.1012 | 8.0877 | 15.28 | 277.2157 | 4.9268 | 17.7 | 594.2796 | 6.2965 |
| 5.19 | 611.1606 | 7.3546 | 11.14 | 271.0597 | 7.5069 | 10.14 | 430.1704 | 4.841 | 14.51 | 277.2157 | 6.038 |
| 15.62 | 175.1478 | 7.2289 | 5.19 | 611.1606 | 6.7041 | 9.97 | 195.1017 | 4.7357 | 14.8 | 411.743 | 5.6639 |
| 12.81 | 387.1792 | 7.0728 | 13.48 | 273.2572 | 6.5637 | 14.8 | 411.743 | 4.6362 | 13.77 | 455.315 | 5.2568 |
| 12.77 | 369.169 | 6.538 | 6.03 | 595.1657 | 6.3949 | 5.19 | 465.1023 | 4.414 | 11.58 | 517.3151 | 5.2171 |
| 17.25 | 609.2706 | 5.8799 | 12.77 | 203.1795 | 6.3935 | 10.14 | 377.1226 | 4.4057 | 11.66 | 517.3153 | 5.2134 |
| 16.61 | 607.2551 | 5.7667 | 6.67 | 303.0494 | 6.182 | 5.53 | 147.0441 | 4.2469 | 13.18 | 453.3355 | 5.1941 |
| 13 | 203.1794 | 5.6487 | 16.16 | 149.1325 | 6.125 | 4.73 | 379.1743 | 4.2169 | 12.64 | 1037.656 | 5.1702 |
| 6.4 | 223.06 | 5.6346 | 6.27 | 625.176 | 6.0964 | 5.19 | 303.0495 | 4.0502 | 10.24 | 303.0491 | 5.0163 |
| 5.47 | 303.0494 | 5.6183 | 16.61 | 607.255 | 5.8769 | 15.02 | 335.2786 | 4.044 | 17.39 | 104.0707 | 4.9004 |
| 5.58 | 303.0494 | 5.5919 | 16.15 | 409.3811 | 5.8144 | 3.66 | 579.1494 | 3.9004 | 14.8 | 784.533 | 4.7003 |
| 6.67 | 303.0494 | 5.5054 | 4.32 | 865.1972 | 5.7156 | 6.27 | 625.176 | 3.8899 | 14.22 | 457.3304 | 4.695 |

**Table S10:** continued

| *Daphniphyllum oldhamii* | | | | | | *Schima superba* | | | | | |
|---|---|---|---|---|---|---|---|---|---|---|---|
| Local Diversity | | | Plot Diversity | | | Local Diversity | | | Plot Diversity | | |
| RT in min | m/z | Comp. 1 | RT in min | m/z | Comp. 1 | RT in min | m/z | Comp. 1 | RT in min | m/z | Comp. 1 |
| 0.99 | 163.0602 | 9.6724 | 17.7 | 593.276 | 16.75 | 1.02 | 193.0708 | 8.5319 | 17.39 | 414.3571 | 10.976 |
| 0.99 | 145.0495 | 8.8168 | 17.7 | 594.2796 | 12.613 | 17.39 | 414.3571 | 8.4745 | 5.19 | 611.1606 | 8.6546 |
| 17.39 | 414.3571 | 6.4792 | 17.46 | 609.2708 | 9.5391 | 5.19 | 611.1606 | 6.9678 | 5.47 | 303.0494 | 7.6712 |
| 14.76 | 200.2009 | 6.4319 | 16.61 | 607.255 | 8.876 | 6.03 | 595.1657 | 6.396 | 6.67 | 303.0494 | 6.679 |
| 17.77 | 352.3387 | 5.7885 | 17.25 | 609.2706 | 8.1555 | 4.28 | 149.0597 | 6.308 | 1.02 | 193.0708 | 6.0124 |
| 10.24 | 303.0491 | 5.6772 | 15.85 | 693.3273 | 7.4269 | 6.67 | 303.0494 | 6.2504 | 5.47 | 465.1023 | 5.9157 |
| 10.22 | 287.0545 | 5.6176 | 0.93 | 266.1594 | 6.9134 | 5.58 | 303.0494 | 6.162 | 5.58 | 303.0494 | 5.2185 |
| 5.66 | 595.1657 | 5.2568 | 6.97 | 433.1121 | 6.809 | 8.08 | 345.169 | 5.4987 | 5.48 | 484.0759 | 5.2074 |
| 5.58 | 303.0494 | 5.2551 | 7.89 | 519.1492 | 6.4785 | 5.47 | 303.0494 | 5.2346 | 6.15 | 303.0493 | 5.1925 |
| 7.89 | 519.1492 | 5.0004 | 11.14 | 271.0597 | 6.1892 | 9.81 | 887.2605 | 5.0333 | 1.03 | 137.0807 | 5.1855 |
| 0.98 | 180.0869 | 4.9711 | 10.22 | 287.0545 | 5.9583 | 10.17 | 873.2443 | 4.9037 | 10.16 | 873.2443 | 5.0684 |
| 12.82 | 255.0647 | 4.9388 | 7.89 | 357.0962 | 5.821 | 12.72 | 1203.615 | 4.7987 | 5.19 | 465.1023 | 4.8255 |
| 16.61 | 607.2551 | 4.8968 | 5.68 | 449.1072 | 5.736 | 14.86 | 457.3662 | 4.7793 | 5.19 | 303.0495 | 4.5264 |
| 14.91 | 98.98411 | 4.8206 | 5.58 | 303.0494 | 5.5867 | 17.65 | 621.2707 | 4.7425 | 17.65 | 621.2707 | 4.4791 |
| 7.89 | 357.0962 | 4.6546 | 4.2 | 175.0392 | 5.419 | 12.29 | 1219.611 | 4.7037 | 10.34 | 873.2443 | 4.4366 |
| 5.68 | 449.1072 | 4.6186 | 16.19 | 231.2106 | 5.0635 | 4.95 | 315.0705 | 4.6522 | 17.18 | 439.3572 | 4.3489 |
| 11.28 | 287.0545 | 4.5845 | 17.43 | 439.3569 | 4.9767 | 5.58 | 465.1022 | 4.633 | 17.7 | 593.276 | 4.3474 |
| 17.22 | 429.0877 | 4.5619 | 7.89 | 147.0441 | 4.6449 | 1.03 | 147.0653 | 4.5234 | 10 | 873.2447 | 4.3339 |
| 4.81 | 597.1446 | 4.5516 | 14.44 | 277.2157 | 4.5933 | 10.25 | 887.2604 | 4.4473 | 11.38 | 439.356 | 4.2329 |
| 6.97 | 433.1121 | 4.4066 | 7.14 | 433.1122 | 4.5903 | 4.29 | 329.1228 | 4.4339 | 17.33 | 483.384 | 4.2161 |

**R Script:**

```
#### Get PubChem CID ####

# install libraries

# load libraries

library(webchem)

library(reshape2)

#---------------------------------#

#read in metaboscape output

#---------------------------------#

dat <- read.csv(file.choose(), sep=",", header = T, stringsAsFactors = F, check.names = F)

# ------------------------------- #

# Get CIDs from molecule names ; credits to Linnea Catherine Smith for solving the curl issue

# ------------------------------- #

# Do it in chunks of 50 because for some reason when running the whole thing at once it gives the error:

# "Error in curl::curl_fetch_memory(url, handle = handle) :

#  Error in the HTTP2 framing layer"  but in smaller batches that doesn't happen

cid.from.name <- get_cid(dat$Name[1:50], from = "name", match = "first", verbose = T)

cid.from.name <- rbind(cid.from.name,get_cid(dat$Name[51:100], from = "name", match = "first", verbose = T))

for(i in seq(nrow(cid.from.name)+1,nrow(dat),by=50)){

  cid.from.name <- rbind(cid.from.name,get_cid(dat$Name[i:(i+49)], from = "name", match = "first", verbose = T))
```

```
  print(paste(i,"/",NROW(dat$Name),sep=""))
}
cid.from.name <- cid.from.name[1:nrow(dat),]
dat$CID.fromName <- cid.from.name$cid
sapply(dat, class)
write.csv(dat,"path/to/your/data.csv",row.names=F)
# ------------------------ #
# get inchikey and smiles from pubchem
# ------------------------ #
# read in or directly use data from previous step
pubchemlist <- dat$cid
pubchemlist <- trimws(pubchemlist)
prop_list <- pc_prop(pubchemlist, properties = c("InchiKey", "Inchi" ,"CanonicalSMILES"),verbose=T)
write.csv(prop_list, "path/to/your/data.csv",row.names=F)
############# Classyfire #################
# read in from previous step
library(RAMClustR)
smiles_info <- getSmilesInchi(dat) # data table from previous step
# get rid of the list history if applicable
input <- smiles_info[-7]
classyfired <- getClassyFire(input, get.all = T, max.wait = 10, posts.per.minute = 5)
```

```
results <- subset(classyfired, select=c("cmpd.name", "CID", "inchikey", "smiles", "classyfire"))

#results[is.na(results)] <- 0

setwd(choose.dir())

write.csv(results, "path/to/your/data.csv",row.names=F)

# merging of classyfire output with original data table with merge function

# sum up intensities per group, done by hand

############ setup data for sunburst plots ###################

#install the packages if necessary

if(!require("tidyverse")) install.packages("tidyverse")

if(!require("fs")) install.packages("fs")

if(!require("readxl")) install.packages("readxl")

if(!require("writexl")) install.packages("writexl")

#load packages

library(tidyverse)

library(fs)

library(readxl)

library(writexl)

path <- choose.files()

mad <- path %>%

  excel_sheets() %>%
```

```
  set_names() %>%
  map(read_excel,
      path = path)
# ---------Plant species used Cgla = Cyclobalanopsis glauca--------------- #
roots.sum <- aggregate(mad$`Cgla root`$`Summed up intensities`,
                       by=list(classyfire.kingdom=mad$`Cgla root`$classyfire.kingdom,
                               classyfire.superclass=mad$`Cgla root`$classyfire.superclass,
                               classyfire.class=mad$`Cgla root`$classyfire.class), FUN=sum)
leaves.sum <- aggregate(mad$`Cgla leaf`$`Summed up intensities`,
                        by=list(classyfire.kingdom=mad$`Cgla leaf`$classyfire.kingdom,
                                classyfire.superclass=mad$`Cgla leaf`$classyfire.superclass,
                                classyfire.class=mad$`Cgla leaf`$classyfire.class), FUN=sum)
exudate.sum <- aggregate(mad$`Cgla exudate`$`Summed up intensities`,
                         by=list(classyfire.kingdom=mad$`Cgla exudate`$classyfire.kingdom,
                                 classyfire.superclass=mad$`Cgla exudate`$classyfire.superclass,
                                 classyfire.class=mad$`Cgla exudate`$classyfire.class), FUN=sum)
x <- list(exudate.sum, leaves.sum, roots.sum)
write_xlsx(x, path = "path/to/your/data.xlsx", col_names=T)
```
